# Molecular Aggregation Induced Photoacoustics for NIR-II in vivo Imaging

**DOI:** 10.1101/2022.04.25.489363

**Authors:** Zong Chang, Liangjian Liu, Chenchen Liu, Shubi Zhao, Jiaqi Chen, Wenxin Zhang, Xiao Wang, Chengbo Liu, Xiaojiang Xie, Qinchao Sun

**Affiliations:** Guangdong Provincial Key Laboratory of Biomedical Optical Imaging Technology & Center for Biomedical Optics and Molecular Imaging, Shenzhen Institute of Advanced Technology, Chinese Academy of Sciences, Shenzhen 518055, China; Department of Chemistry, Southern University of Science and Technology, Shenzhen, 518055, China; Shenzhen Key Laboratory of Nanobiomechanics, Shenzhen Institute of Advanced Technology, Chinese Academy of Sciences, Shenzhen 518055, China; Shenzhen College of Advanced Technology, University of Chinese Academy of Sciences, Shenzhen 518055, People’s Republic of China; National Innovation Center for Advanced Medical Devices, Shenzhen, 518131, People’s Republic of China

## Abstract

Molecular aggregation induced photo-properties alteration has been found to play a crucial role in the light induced processes, such as aggregation induced emission (AIE) and J aggregation induced dramatic absorption red shift. The light induced acoustic process (photoacoustic) is also considered to be one of the most essential characters of the light absorbing molecules. However, to the best of our knowledge, the molecular aggregation induced photoacoustic effect (MAIPA) has never been reported. Herein, we report the first MAIPA effect for which the PA intensity is dominated by the molecular aggregation, rather than by absorbance as usual concerned. Molecular aggregation induces a strong electronic coupling effect, resulting in significant absorption suppression from the individual state to highly aggregated state (around 5 molecules aggregated). However, the corresponding PA efficiency was found to be about 2-orders of magnitude greater for the latter. A well-behaved linear correlation between the molecular aggregation level and MAIPA effect was observed. The surprisingly significant MAIPA effect was realized via novel NIR-II squaraine-benzothiopyrylium dyes. Excellent photophysical properties of the novel NIR-II dyes were achieved, such as large absorption extinction coefficient and high photostability. Thanks to the relatively narrow FWHM and the high PA efficiency of SQN2@PMAOPEG and ZC825@BSA, in vivo multiplex PA imaging was demonstrated for tumor tissue and macrophage cells, blood and lymphoid vessels.

## Introduction

Molecular aggregation induced photo-properties alteration has been intensively studied for centuries, which plays a crucial role not only in the artificial system but also in nature.^1,2^ For instance, the J aggregated bacteriochlorophylls of the light harvesting center (LHC) would render the photosynthetic bacteria utilising the low energy sunlight for biochemical reaction.^1^ Furthermore, the molecular aggregation induced emission (known as AIE) has been recently discovered and widely used for biological fluorescence imaging, diagnosis, etc.^2,3^ Regarding to the AIE effect, the molecules are in a confined space to inhibit the nonradiative decay processes (such as molecular rotation) to enhance the fluorescence emission efficiency. Besides the light induced emission and chemical reaction processes, the light induced acoustic (photoacoustic) is also considered to be one of the most essential characters of the light absorbing molecules.^4–6^ Though, a few studies have found aggregation related PA enhancement, for instance, in the case of J aggregation induced new absorption band,^7^ the formation of the large size cluster of nanoparticles or molecules.^8,9^ To the best of our knowledge, the molecular aggregation induced photoacoustic effect (MAIPA) for which decoupling from the absorbance and size increasement effect has never been reported.

Photoacoustic (PA) imaging as one of the most substantial imaging techniques has been studied intensively over the last decades.^4–6^ Taking the advantage of the optical excitation and the ultrasound propagation properties, the PA imaging provides much higher image contrast than ultrasound imaging (only the region with the right light absorber could be imaged), and much deeper tissue penetration than optical imaging (much lower tissue scattering for ultrasound than light). Therefore, PA imaging exhibits great potential for clinical medicine and in vivo biomedical research.^6,10^ To achieve high-performance PA imaging, the light radiation should be efficiently converted to acoustic waves via thermal expansion. The light absorber (contrast agent) as the energy converter is considered to be the most critical role for PA imaging. Besides a few endogenous contrast agents in specific cells, for instance, the haemoglobin in blood cells is able to provide clear and informatic images of blood vessels, most of the cells are lack appreciate light absorbing agents.^11^ Therefore, artificial contrast agents are necessary to realize the in vivo PA imaging for more general applications. In recent years, an enormous amount of photoacoustic contrast agents has been developed, for instance, small organic molecules,^12^ metallic nanoparticles,^13^ semiconducting polymer,^14^ inorganic materials,^15^ etc.^16,17^ Furthermore, the NIR-II PA contrast agents have attracted more and more attention.^18–20^ The relative lower tissue scattering loss of the NIR-II excitation beam was considered to further improve the penetration depth for PA imaging.^20^

Organic dyes are widely applied for in vivo PA and fluorescence imaging, due to the relatively low cytotoxicity and great clinical translation potential.^21^ Nevertheless, as the absorption energy shifting to NIR-II region, the photostability and absorption extinction coefficient would be a great challenge for organic dyes. Furthermore, the dye micelle system is one of the mostly taken strategy for in vivo administration of PA contrast agent to improve the biocompatibility and to prolong the blood circulation time, especially for NIR-II contrast agents.^22,23^ In general, to retain the photophysical properties of dyes, the mass ratio between dye and amphiphilic molecules was applied as large as possible.^23^ However, the amphiphilic molecules (micelle) are known to be highly toxic to damage the cell membrane (e.g. hemolysis), therefore, the amount of which applied for in vivo study should be carefully controlled (the lower the better).^24^ The administration amount of the dyes micelle system would be compromised between the in vivo tolerance of the cytotoxicity and in vivo PA signal. Furthermore, currently developed PA imaging contrast agents provide outstanding images of blood vessels, tumor tissue, etc. However, most of the reported PA contrast agents were used as standalone agents for mono-tissue imaging.^25,26^ To develop novel contrast agent pairs for multiplex PA imaging would greatly advance our understanding of in vivo biomedical processes, for instance, the interaction between the tumor and immune cells during immunotherapy.^27^

A series of novel NIR-II squaraine-benzothiopyrylium dyes were designed with high absorption extinction coefficient and photostability to address abovementioned limitations. The corresponding absorption transition energy was able to be gradually adjusted via varying the electron donation groups on benzothiopyrylium. Thanks to the relatively narrow FWHM of SQN2 (maximum at 1085 nm) and ZC825 (maximum at 825 nm), a high contrast multiplex PA imaging experiments was able to be realized with excitation at 1064 nm and 825 nm. A surprisingly significant MAIPA effect was observed via squaraine-benzothiopyrylium dyes. In general, the contrast agent with higher absorption cross-section is necessary to gain the greater PA signal. Herein, we found that the molecular aggregation dominates the PA intensity over the absorbance. In the case of dye SQN2, molecular aggregation induces a strong electronic coupling, resulting in significant absorption suppression as much as 8-folds from the individual state to highly aggregated state (around 5 molecules). However, the PA efficiency was found to be about 2-orders of magnitude greater for the highly aggregated state than the individual state. The MAIPA effect would open a new window for the design of PA contrast agent systems, for instance low cytotoxicity theranostic micelle system and low dose light irradiation PA imaging.

## Results and Discussion

### Photophysical properties characterization

The synthesized cyanine-like and squaraine-benzothiopyrylium dyes were shown in Figure 1a and S1. Via rational design of the electron properties of the substitution group on benzothiopyrylium and the corresponding bridge chain, the absorption maximum could be gradually red shifted from 1003 nm to 1085 nm, as shown in Figure 1b. For CyNPh, CyNO and CyN2, the benzothiopyrylium groups are linked by a pentamethine chain to form a cyanine-like electronic structure. The electronic donation property was gradually enhanced from phenylamine to alkylamine and then to dual alkylamine groups for CyNPh, CyNO and CyN2 respectively. The corresponding absorption maximum was found to be 1003 nm, 1020 nm and 1031 nm. A similar electronic effect was found in the case of squaraine bridge-linked benzothiopyrylium dye, with about 30 nm red shift from SQNPh to SQN2. In comparison to the pentamethine bridge chain, the squaraine bridge would further lower the absorption energy from 1031 nm to 1085 nm with the same electron donation group of benzothiopyrylium for CyN2 and SQN2. The HOMO and LUMO molecular orbital of the synthesized dyes were shown in Figure 1c and Figure S2. The HOMO and LUMO electrons were highly delocalized on the benzothiopyrylium and the bridge chain (pentamethine, squaraine). However, for the squaraine-benzothiopyrylium dyes, a higher electron density on the nitrile group of squaraine was found in the HOMO molecular orbital than that in the LUMO. It indicates that an intramolecular charge transfer (ICT) undergoes from the squaraine group to the main molecules in the excited state. The ICT might induce the large red shift in absorption from CyN2 to SQN2.^21^ The key point to achieve a successful multiplex photoacoustic imaging is the well separated absorption spectra of the contrast agent. The SQN2 with an absorption maximum around 1085 nm and a relatively narrow absorption band (FWHM about 80 nm) exhibits the right photo-properties for multiplex imaging. The ZC825 with maximum absorption around 825 nm was taken for the second contrast agent. From Figure 1d, we could find the unambiguous separation of the absorption spectra of SQN2 and ZC825 at the excitation wavelength about 1064 nm and 825 nm. Furthermore, the SQN2 exhibits strong absorption extinction coefficient and high photostability, as shown in Figure S3 and S4.

**Figure 1.**
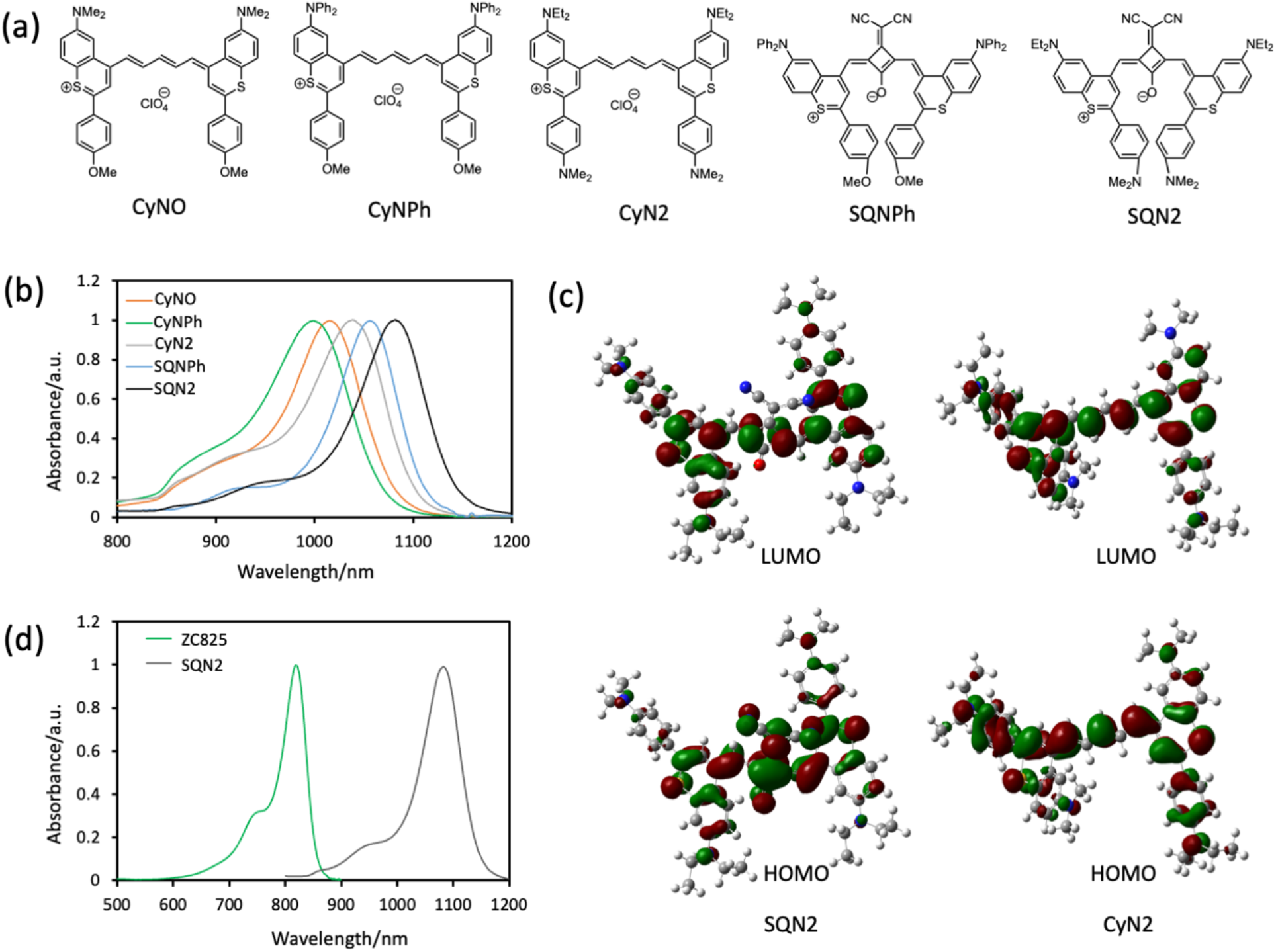
The molecular structure and electronic properties of the synthesized NIR-II dyes. **(a)** The molecular structure of the synthesized NIR-II dyes. **(b)** The absorption spectra of synthesized NIR-II dyes in DCM solution. **(c)** The HOMO and LUMO molecular orbitals of SQN2 and CyN2. **(d)** The normalized absorption spectra of SQN2 in DCM and ZC825 (Figure S1) in EtOH.

### Aggregation of Molecules

To improve the biocompatibility of the prepared PA contrast agent, the BSA (bovine serum albumin) and PMAOPEG micelle were applied. The BSA as the natural nutrition delivery carrier has been widely used to make materials more biocompatible such as nanoparticles and small organic dyes, etc.^28,29^ The BSA was found to be able to efficiently bind with ZC825 molecules. As the mass ratio between ZC825 and BSA varying from 1:200 to 1:88, the absorption spectra of ZC825@BSA remains similar (the mass ratio at 1:88 corresponding to the similar molar quantity of ZC825 and BSA), as shown in Figure 2a. While the mass ratio of ZC825: BSA beyond 1:88, the absorption maximum of ZC825@BSA in aqueous solution was reduced significantly from about 1 (ZC825: BSA at 1:88) to 0.5 (ZC825: BSA at 1:20). The colour change of the ZC825@BSA as a function of mass ratio could be visualized by naked eyes under the same mass concentration of the ZC825, as shown in Figure 2c. The PMAOPEG was synthesized as an amphiphilic polymer to prepare the SQN2@PMAOPEG micelle since the BSA was not able to bind effectively with SQN2. As the mass ratio of SQN2 to PMAOPEG varying from 1:1000 to 1:20, the absorbance of SQN2@PMAOPEG in aqueous solution was dramatically suppressed by a factor of around 8, from OD about 2.0 to 0.25 at the absorption maximum. The reduction of the absorption intensity at high mass ratio of ZC825@BSA and SQN2@PMAOPEG might be caused by the strong electronic coupling via aggregation of molecules. For the case of high amounts of BSA or PMAOPEG, the absorption spectrum of ZC825@BSA (from 1:200 to 1:88) and SQN2@PMAOPEG (1:1000) in aqueous was found to be similar to that in organic solvent, Figure 1d, Figure 2a, 2b, and Figure S5. Regarding to the absorption peak to shoulder ratio, it was about 4 for SQN2@PMAOPEG in aqueous solution (about 5 for SQN2 in DCM), and about 3 for both ZC825@BSA and ZC825 in EtOH, as shown in Figure S5. It indicates the individual molecule was investigated for ZC825@BSA (from 1:200 to 1: 88) and SQN2@PMAOPEG (1:1000).

**Figure 2.**
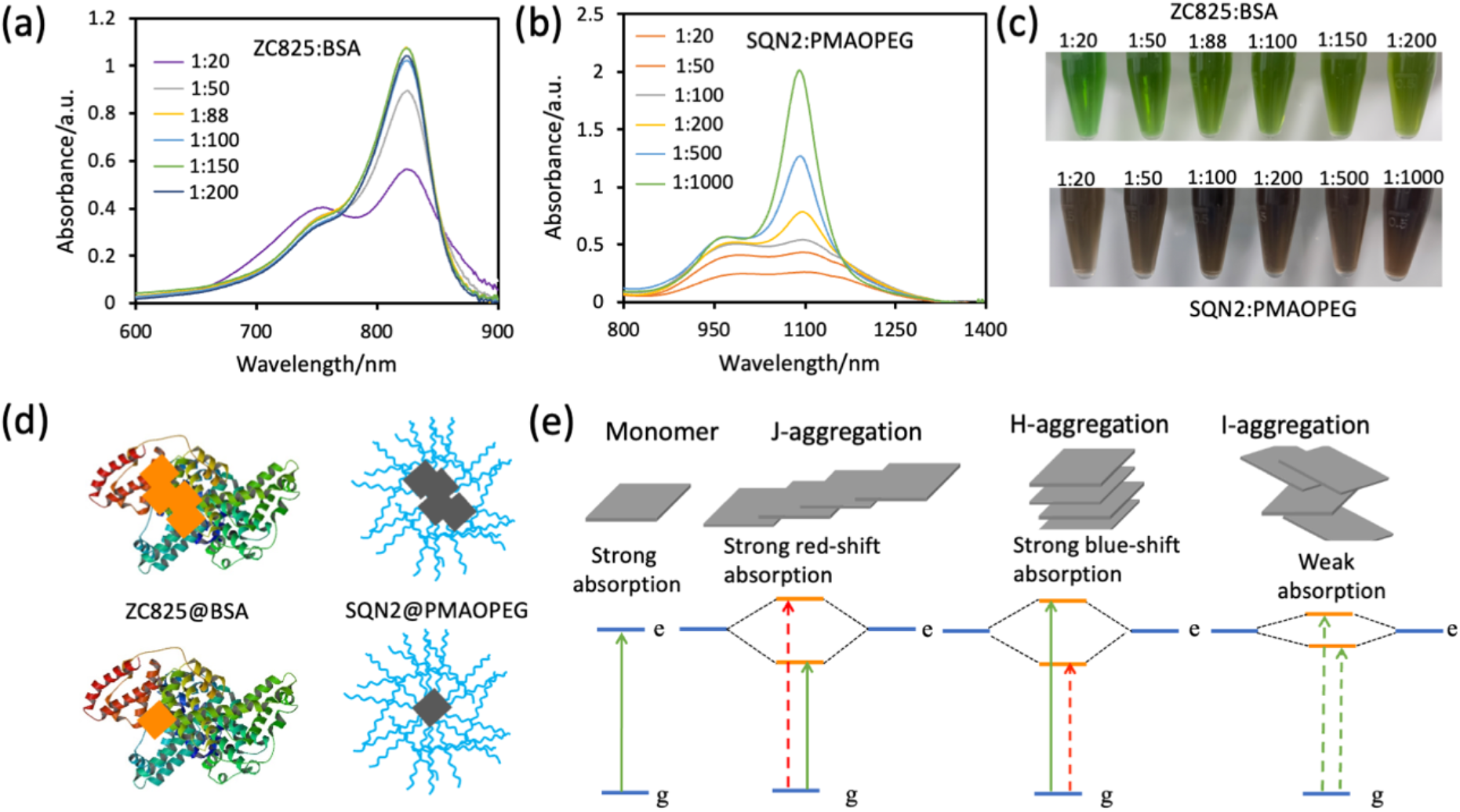
The photophysical properties of the ZC825@BSA and SQN2@PMAOPEG. **(a)** The absorption spectra of the ZC825@BSA as a function of the mass ratio of ZC825 to BSA from 1:20 to 1:200 in aqueous solution with ZC825 mass concentration constant (25 μg/mL). **(b)** The absorption spectra of SQN2@PMAOPEG as a function of the mass ratio of SQN2 to PMAOPEG from 1:20 to 1:1000 in aqueous solution with SQN2 mass concentration constant (50 μg/mL). **(c)** The white light image of the prepared ZC825@BSA and SQN2@PMAOPEG. **(d)** The scheme of the molecular aggregation of ZC825 and SQN2 interacting with BSA and PMAOPEG, respectively. **(e)** The scheme of molecular aggregation induced electronic properties variation; the I-aggregation represents the case that the molecules do not aggregate in the ideal H/J structure (nonideal aggregation).

For ZC825@BSA at mass ratio of 1:88, the individual ZC825 binding with single BSA was consistent with the corresponding molar ratio (1:1), due to the specific interaction between each other. However, for the case of SQN2@PMAOPEG micelle, the interaction between each other is more random. The number of SQN2 in single PMAOPEG micelle could be addressed by the Poisson distribution as described in the methods section. The number of PMAOPEG micelles should be around 10 times more than SQN2 to realize individual SQN2 molecule involving in single PMAOPEG micelle as mass ratio about 1:1000, which is coincident as revealed by the absorption spectrum, as mentioned above as individual state (Figure S5). Furthermore, the average number of SQN2 molecules in single PMAOPEG micelle at mass ratio about 1:20 was about 5. The average number of ZC825 on single BSA at mass ratio 1:20 was about 4, according to the corresponding molar ratio.

The lower amount of the BSA or PMAOPEG would result in more than one dye molecule interacting with a single BSA, or PMAOPEG micelle as illustrated in Figure 2d. The molecules in such case would aggregate tightly to induce a strong electronic coupling. The well-known J/H aggregation would introduce strong electronic coupling leading to a red/blue shifted absorption in comparison to the individual molecule, as illustrated in Figure 2e.^30,31^ In most of the cases, the aggregated molecules orient to each other neither in the ideal H nor J geometry. Nevertheless, the electronic coupling between each other remains significant. Consequently, the electronic transition dipole moment of such tightly aggregated molecules would be dramatically reduced in comparison to the individual molecule. The corresponding absorption intensity would be suppressed (without significant blue/red shifted absorption), which might give a reasonable explanation of the photophysical properties variation of SQN2@PMAOPEG and ZC825@BSA as the mass ratio changes.

### Molecular aggregation induced PA

It is well-known, the PA intensity is proportional to the thermal conversion of the light absorber and the Grüneisen parameter of the system, as shown in Equation (1).

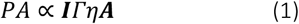

Where ***I*** is the excitation beam intensity, *Γ* is the Grüneisen parameter which reveals the thermal expansion of the system, the *η* is the nonradiative decay efficiency of light absorbers and **A** is the absorbance.

The thermal energy conversion efficiency *η* of ZC825@BSA and SQN2@PMAOPEG in aqueous solution was considered to be close to unity. The emission spectra of ZC825@BSA and SQN2@PMAOPEG were shown in Figure S6. Even though a clear emission spectrum could be recorded for both ZC825@BSA and SQN2@PMAOPEG in aqueous solution, the emission quantum yield of ZC825@BSA is found to be no more than 1% and for SQN2@PMAOPEG less than 0.1%. From Eq (1), we may find that with the same thermal expansion properties (Grüneisen parameter) of a system, the higher the absorbance of the light absorber the larger the PA intensity. The PA intensity of ZC825@BSA and SQN2@PMAOPEG was measured in aqueous solution as shown in Figure 3a, 3b. We surprisingly found that the lower the absorbance of the aggregated light absorber the higher the PA intensity was recorded. For ZC825@BSA, as the absorbance decreased from 1 to 0.5, from the individual state (mass ratio 1:200) to the highly aggregated state (mass ratio 1:20), the PA intensity was found to be increased by about 3-fold (Figure 3a, same mass concentration of ZC825). For SQN2@PMAOPEG, under the same mass concentration, the absorbance of the highly aggregated state (mass ratio 1:20) was suppressed by a factor of 5 in comparison to that of the individual state (mass ratio 1:1000) at 1064 nm. However, the PA intensity of the highly aggregated state (around 5 molecules aggregated) was about 30-fold higher than the individual state, as shown in Figure 3b.

**Figure 3.**
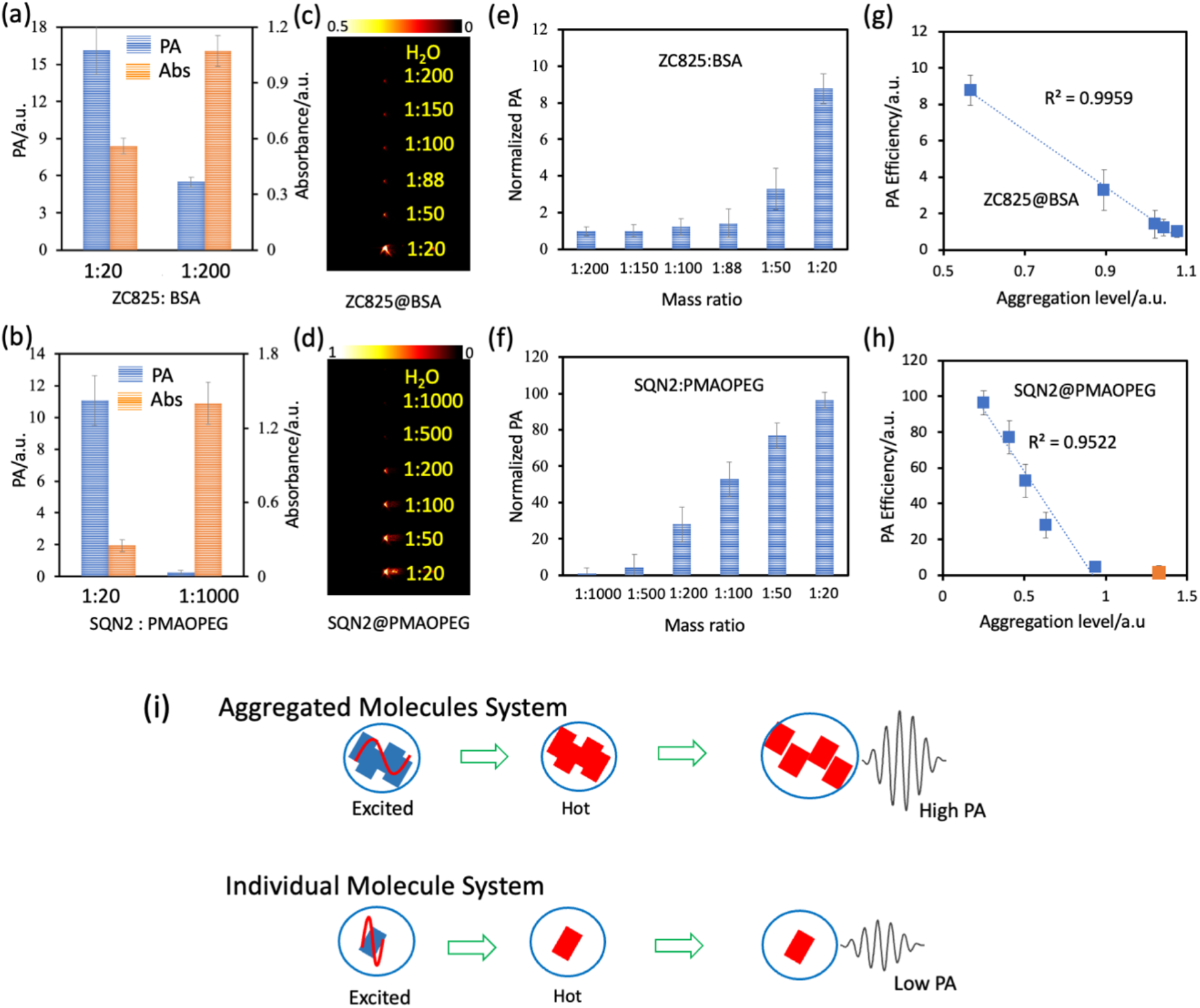
The photoacoustic properties of the ZC825@BSA and SQN2@PMAOPEG. **(a)** The PA intensity of ZC825@BSA excited at 825 nm (left axis), and absorbance at 825 nm (right axis) at mass ratio of 1:20 and 1:200, (ZC825, 25 μg/mL). **(b)** The PA intensity of SQN2@PMAOPEG excited at 1064 nm (left axis), and absorbance at 1064 nm (right axis) at mass ratio of 1:20 and 1:1000 (SQN2, 25 μg/mL). **(c)** The PA image of ZC825@BSA with a mass ratio from 1:20 to 1:200 with the same absorbance about 1 at 825 nm with ZC825 concentration about 25 μg/mL for mass ratio (1:200 to 1:88), 29 μg/mL for 1:50 and 34 μg/mL for 1:20. **(d)** The PA image of SQN2@PMAOPEG with the same absorbance about 1 at 1064 nm with SQN2 concentration about 22, 42, 67, 103, 128, 209 μg/mL, for mass ratio from 1:1000 to 1:20. **(e)** The normalized PA intensity of ZC825@BSA at mass ratio from 1:20 to 1:200 with the same absorbance about 1 at 825 nm, the PA intensity at mass ratio 1:200 as 1. **(f)** The normalized PA intensity of SQN2@PMAOPEG at mass ratio from 1:20 to 1:1000 with the same absorbance about 1 at 1064 nm, the PA intensity at mass ratio at 1:1000 as 1. **(g)** The PA efficiency of ZC825@BSA as a function of aggregation level (square) and the corresponding linear fitting function (dotted line). **(h)** The PA efficiency of SQN2@PMAO as a function of aggregation level (square) and the corresponding linear fitting function (dotted line), the orange square represents the data at mass ratio of 1:1000. **(i)** The scheme of the hypothetical mechanism of the molecular aggregation induced photoacoustics (MAIPA).

Figure 3c and 3d show the PA image of ZC825@BSA and SQN2@PMAOPEG at the excitation wavelength about 825 nm and 1064 nm respectively, under the absorbance of about 1 (1 cm optical path, at excitation wavelength). From Eq (1), we might define the PA efficiency of a system by *PA*/***AI***, which would be constant for a certain system. Figure 3e and 3f depict the relative PA efficiency with the same absorbance at different mass ratios for ZC825@BSA and SQN2@PMAOPEG. The PA efficiency of ZC825@BSA and SQN2@PMAOPEG was normalized to the corresponding individual state. It was found that the PA efficiency of ZC825@BSA remains similar for the mass ratio from 1:200 to 1:88. It is in accordance with the aggregation level of the ZC825@BSA system, as shown in Figure 2a, where the variation on absorbance from 1:200 to 1:88 was also negligible. Whereas the PA efficiency increased dramatically by about 10-fold from the mass ratio of 1:88 to 1:20. For the case of the SQN2@PMAOPEG system, under the same absorbance of the excitation beam, the PA efficiency increased gradually as the corresponding mass ratio became larger. The PA efficiency was found to be about 2-orders of magnitude higher for highly aggregated states (1:20) than for the individual state (1:1000).

To study the relationship between the PA efficiency and the aggregation level, the PA efficiency as a function of the aggregation level was plotted in Figure 3g and 3h. The aggregation level was evaluated by the absorbance at different mass ratios for ZC825@BSA and SQN2@PMAOPEG, since the absorbance is directly affected by the aggregation of molecules (Figure 2a, 2b). For ZC825@BSA, a highly linear correlation between the PA efficiency and the aggregation level was found as the R square close to 1. For SQN2@PMAOPEG, the PA efficiency was also found to be significantly linearly correlated to the aggregation level with R square about 0.95 for mass ratio from 1: 500 to 1: 20. The PA efficiency at the mass ratio of 1:1000 fell off the linear fitting curve. It might come from the fact that the corresponding PA intensity of which is in the similar range of water background.

The well-behaved linear correlation between the PA efficiency and the aggregation level provides further evidence that the molecular aggregation would induce an enormous PA signal. It is well-known that strong electronic coupled aggregated molecules act as a macromolecule, which shares the same electronic state. The aggregated molecules reach the excited state simultaneously while absorbing a photon, as illustrated in Figure 3i. The excited aggregated molecules would simultaneously convert the electronic energy to thermal energy via internal conversion. The simultaneously heated aggregated molecules probably undergo an intra-macromolecule expansion between each other, which might be the origin of such huge PA signal. From Eq (1), we find that the PA efficiency is proportional to the capability of thermal expansion (Grüneisen parameter). While the existence of BSA or PMAOPEG in solution probably make the total Grüneisen parameter of the system a bit higher, as most of the matters have larger Grüneisen parameter than water.^32^ However, we found that such kind of effect plays negligible role in our research, since the corresponding mass concentration less than 2.5% (for instance, in the case of ZC825@BSA from 1:200 to 1:88, the BSA concentration reduced about 2-fold, while no significant PA signal variation observed, Figure 3e.), and the PA signal getting higher as the BSA or PMAOPEG concentration getting lower. Furthermore, there was no significant variation observed on the hydrodynamic size of the ZC825@BSA (around 10 nm) and SQN2@PMAOPEG (around 100 nm) at different mass ratio as shown in Figure S7, which ruled out the size increasement caused PA enhancement. Therefore, from the observation of the SQN2@PMAOPEG and ZC825@BSA, we may find that the purely aggregated molecules play a dominant role in PA efficiency.

### Multiplex in vivo imaging

The MAIPA effect discovered herein ideally addressed the paradox of the PA efficiency and the cytotoxicity of dye micelles system. To retain high absorbance of dyes would not come into the first place for achieving significant PA signal anymore. In the MAIPA system, lowering the amount of amphiphilic molecules would lead to a much higher the PA signal. Under the same concentration of PMAOPEG, the PA intensity of SQN2@PMAO-PEG (1:1000) was not able to be distinguished from the background as shown in Figure S8. While an enormous PA signal was observed for SQN2@PMAO-PEG (1:20) with the same experimental condition. Taking advantage of the MAIPA effect to reduce the amount of amphiphilic agent, high cell viability and no organ damage were observed. The cytotoxicity of the ZC825@BSA and SQN2@PMAO-PEG was evaluated by the CCK8 and the H&E stain of organs, shown in Figure S9, S10.

Immunotherapy as one of the most promising cancer therapeutic measures has been studied intensively in recent years.^33^ However, the in vivo observation of the interaction between the tumour and immune cells is considered to be a great challenge. Multiplex PA imaging would provide the ideal tool for distinguishing such two kinds of cells in vivo. The SQN2 and ZC825 prepared herein exhibit a fully separated absorption spectrum as shown in Figure 1d. The unambiguous distinction of the PA signal between the SQN2@PMAOPEG and ZC825@BSA (mass ratio 1:20) with excitation wavelengths at 1064 nm and 825 nm was realized, as shown in figure 4a and 4b. The PA ratio for both ZC825@BSA at 825 nm/1064 nm and SQN2@PMAO-PEG at 1064 nm/825 nm was about 20. The Hela cells and macrophage cells (stained with ZC825@BSA and SQN2@PMAOPEG) were subcutaneously injected into a mouse model at different locations. The corresponding PA images with excitation at 825 nm and 1064 nm were shown in Figure 4c. The location of Hela cells and macrophage cells could be clearly distinguished, as shown by the green and red-hot channels. Thanks to the absorption of haemoglobin and water at 1064 nm, the blood vessels could also be visualized coincidentally as presented in the red-hot channel. The PA ratio for Hela cells at 825 nm/1064 nm is about 40, while for macrophage cells that at 1064 nm/825 nm is about 50. The remarkable PA performance of ZC825@BSA and SQN2@PMAOPEG indicates their great potential for in vivo multiplex imaging. We further performed a subcutaneous tumour model, for which the Hela cells were ex vivo stained with ZC825@BSA and then inoculated. When the tumour grew to a diameter about 3 mm, the SQN2@PMAOPEG stained macrophage cells were subcutaneously injected near the location of the tumour. The PA image of the tumour and macrophage cells were shown in Figure 5a and 5b. The relative spatial distribution of the macrophage cells and tumour was reconstructed in Figure 5c. Moreover, the depth dependent PA image was shown in Figure 5d with excitation at 1064 nm. The blood vessels and macrophage cells distribution in the Z direction was able to be observed.

**Figure 4.**
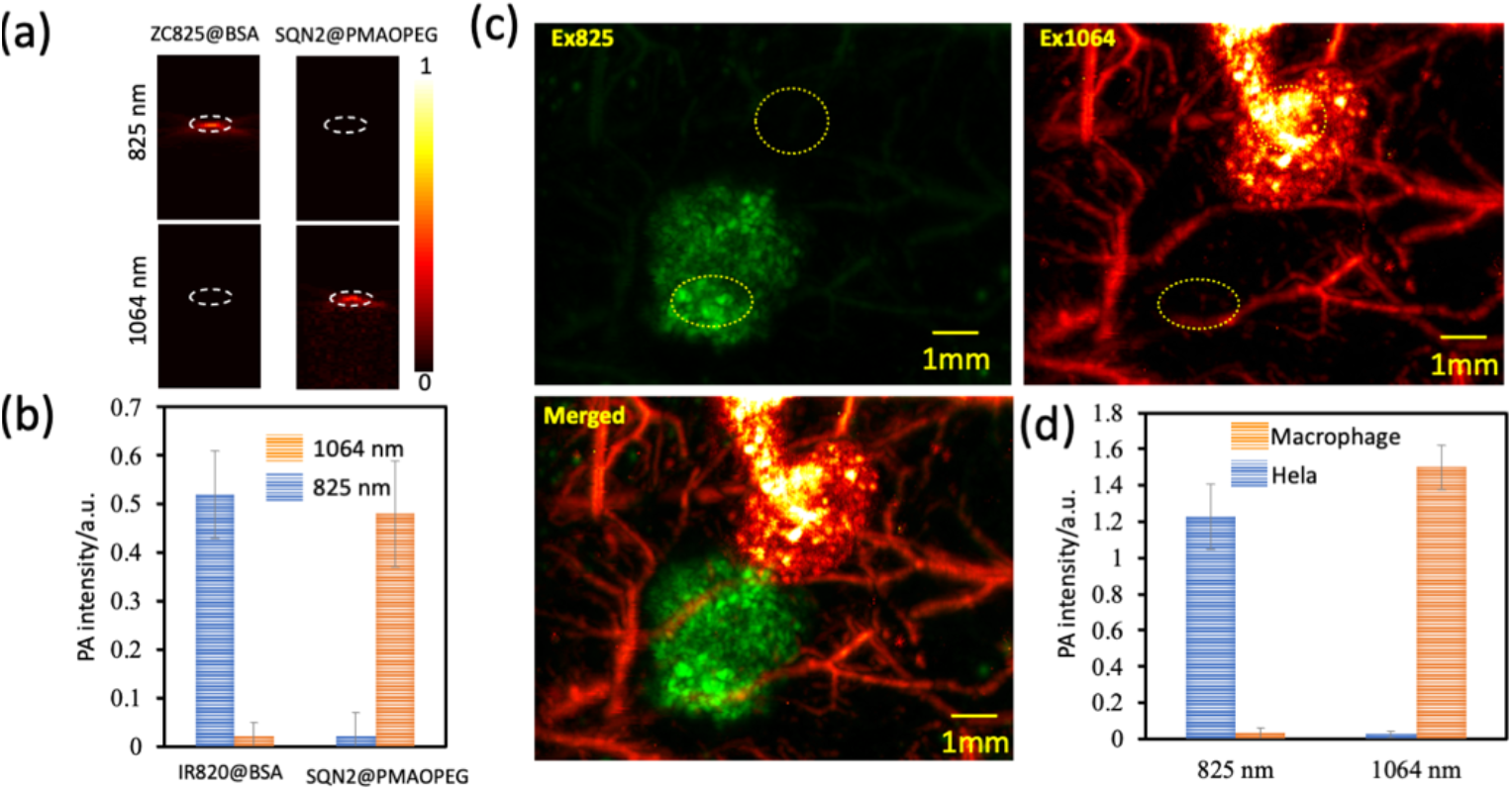
In vivo photoacoustic imaging of ZC825@BSA stained Hela cell and SQN2@PMAOPEG stained macrophage cell. **(a)** The PA image of ZC825@BSA and SQN2@PMAOPEG at mass ratio 1:20 with excitation at 825 nm and 1064 nm. **(b)** The PA intensity of ZC825@BSA and SQN2@PMAOPEG at mass ratio 1:20 with excitation at 825 nm and 1064 nm. **(c)** The PA image of stained Hela cells (green channel) and macrophage cells (red hot channel) via subcutaneous injection with excitation at 825 nm, 3 mJ and 1064 nm, 15 mJ/pulse. **(d)** The quantitative analysis of the PA intensity of ROI at excitation at 825 nm and 1064 nm, respectively, as shown in the yellow circle in (c).

**Figure 5.**
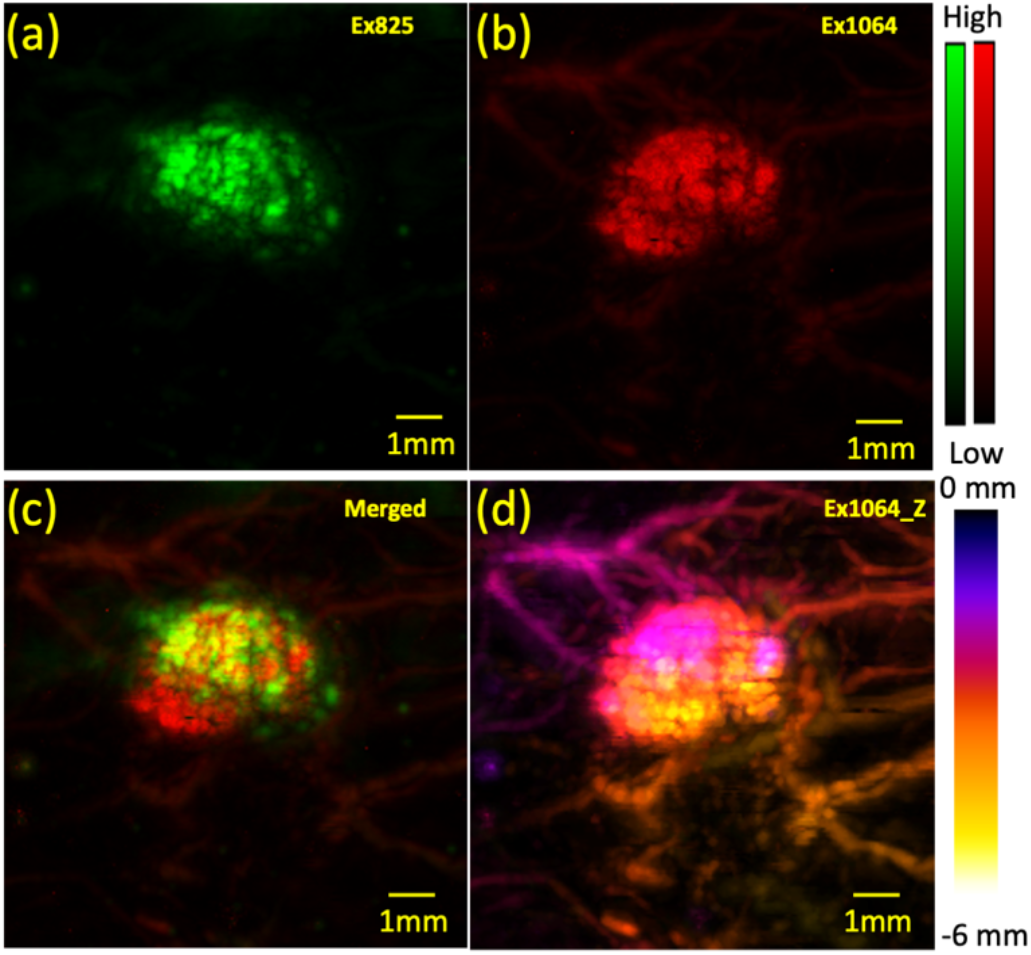
In vivo PA imaging of tumour tissue and macrophage cells. **(a)** The subcutaneous inoculated tumour which was stained by ZC825@BSA before inoculation and then in vivo grew for about a week (excitation at 825 nm, 3 mJ/pulse). **(b)** The stained macrophage cells with SQN2@PMAOPEG via subcutaneous injection under excitation at 1064 nm, 15 mJ/pulse. **(c)** The merged image of the tumour tissue (green) and the macrophage (red). **(d)** The depth dependent color map (Z-location) of the blood vessels and macrophage cells at the excitation of 1064 nm.

The blood and lymphoid circulation system are the two most important transportation systems of mammals. PA imaging on the blood vessels has been widely studied, however to record the clear PA image of lymphoid system is a much more challenge task.^25,34^ The ZC825@BSA was intravenously administered for PA imaging of the blood vessel network, as shown in figure 6b. The lymphoid vessel was visualized by subcutaneous injection of SQN2@PMAOPEG. Figure 6d depicts the relative location of the lymphoid vessels (green channel) and the blood vessels (red channel). Even though the blood vessel could also give a PA signal at 1064 nm, the lymphoid vessel could be clearly distinguished with a much higher signal to background ratio about 4, as shown in Figure 6f. For blood vessel imaging with excitation at 825 nm, a signal to background ratio of about 4 (0.12/0.027) could also be achieved (Figure 6e).

**Figure 6.**
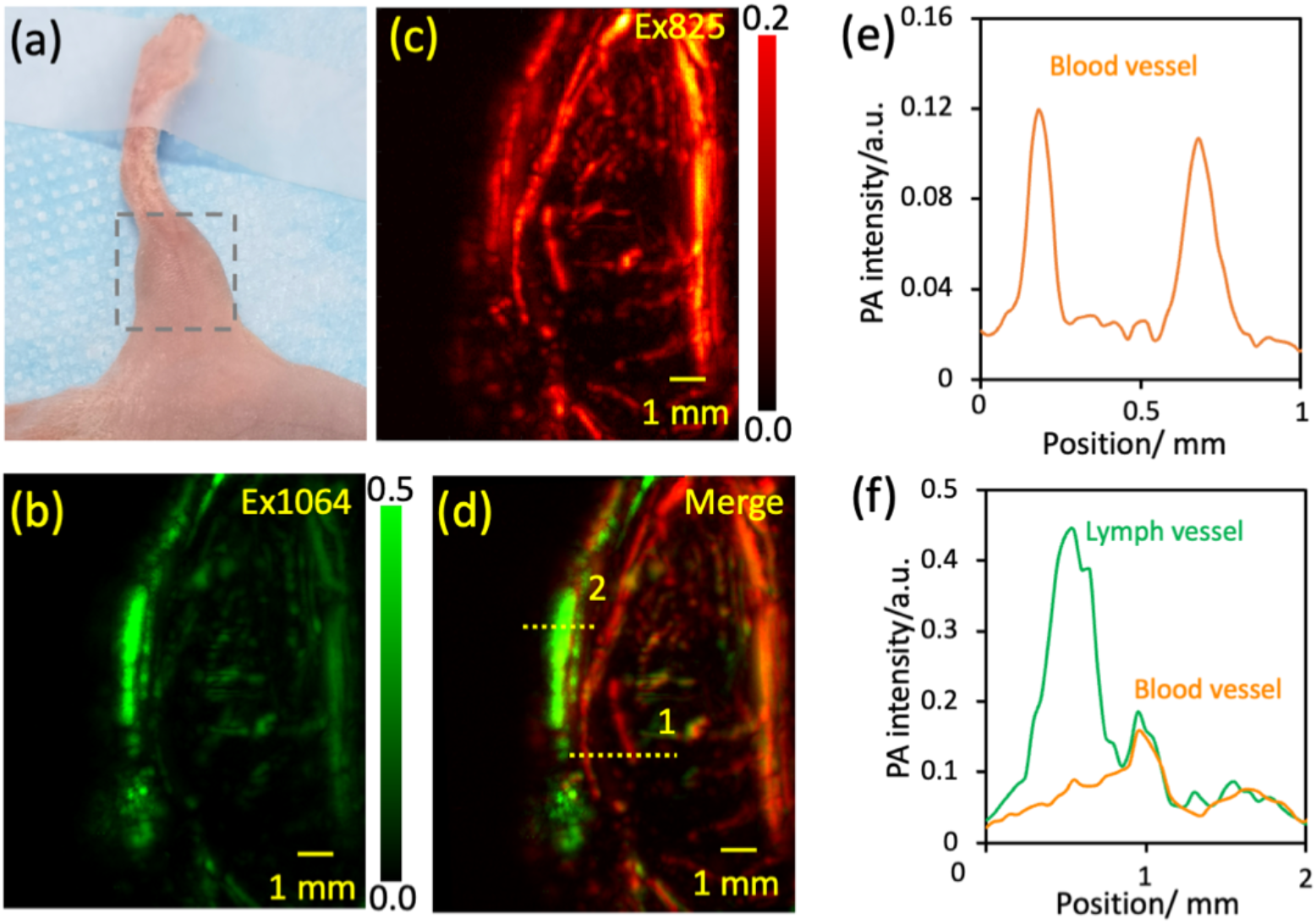
In vivo PA imaging of the blood and lymph vessels. **(a)** The photograph of the region of interest for PA imaging (grey square, left hindlimb). **(b)** The PA image at 30 mins after subcutaneous injection of SQN2@PMAOPEG into the right paw with excitation at 1064 nm, 15 mJ/pulse. **(c)** The PA images of blood vessels via intravenous administration of ZC825@BSA with excitation at 825 nm, 5 mJ/pulse. **(d)** The merged PA images of the lymph vessels (green) and blood vessels (red). **(e)** The PA intensity of the ROI (yellow dotted line (1) in (d) red channel). **(f)** The PA intensity of the ROI in (d) yellow dotted line (2).

## Conclusion

In conclusion, we first reported the molecular aggregation induced photoacoustic (MAIPA) effect, for which the molecular aggregation plays much more dominant role than molecular absorption. The PA efficiency for the highly aggregated state (around 5 molecules aggregated) was observed to be about 2-orders of magnitude greater than the individual state. A well-behaved linear correlation was found between the PA efficiency and the molecular aggregation level. The surprisingly significant MAIPA effect was realized by novel squaraine-benzothiopyrylium NIR-II dye. A series of novel NIR-II dyes have been designed with gradually red shifted absorption from 1003 nm to 1085 nm. The squaraine-benzothiopyrylium NIR-II dye exhibits large NIR-II absorption extinction coefficient and high photostability. Thanks to the fully separated absorption spectrum of SQN2 and ZC825, multiplex PA imaging was performed with excitation at 1064 nm and 825 nm. The great potential of SQN2@PMAOPEG and ZC825@BSA for multiplex in vivo PA imaging was demonstrated by the study of the relative location of tumour cells and the macrophage cells, the blood and lymphoid vessels. The discovery of the MAIPA effect would open a new window for the design and utilization of the PA contrast agent for in vivo imaging, for instance low cytotoxicity micelle theranostic system and low dose light irradiation PA imaging.

## Methods

### Materials

All reagents and solvents were used as received without further purification. Poly (maleic anhydride-alt-1-octadecene) (PMAO), dimethyl sulfoxide (DMSO) and bovine serum albumin (BSA) (98%) were purchased from Sigma-Aldrich (St. Louis, USA). Roswell Park Memorial Institute 1640 (RPMI 1064) Medium and Penicillin–Streptomycin were purchased from HyClone. Phosphate-buffered solution (PBS) was purchased from Corning. Pancreatin was purchased from Coolaber. Dulbecco’s Modified Eagle’s Medium (DMEM) and fetal bovine serum (FBS) were purchased from Gibco. Dojindo Chemical Technology (Shanghai) Co., Ltd supplied Cell Counting Kit-8 (CCK-8) for Cell proliferation and toxicity test. All other chemicals were bought from Aladdin (Shanghai, China) or Energy Chemical (Shanghai, China). Deionized water (18.25 MΩ cm, 25 °C; Millipore, Billerica, USA) was used for all experiments. Flash chromatography was performed on columns of silica gel (300-400 mesh) supplied by yantai Silica Gel Factory (China).

### Number of dyes per micelle

The general preparation procedure for dye embedded in micelle takes place in a quite randomly manner, since there is no specific interaction between each other. The number of dyes per micelle might be described by the Poisson distribution, as followed equation.

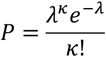

Where *λ* is the average possibility of the number of dyes per micelle, *κ* is the number of dyes per micelle.

In order to make sure a single dye molecule associated in one micelle, the *λ* should be around 0.1. In that case, about 90% micelle with no dye inside. The number of micelles with one dye inside is about 10 times more than the number of micelles with two dyes in. The *λ* around 0.1 corresponds to the molar ratio of dye to micelles about 10. In the case of SQN2@PMAOPEG, The *λ* around 0.1 corresponds to the mass ratio at 1:1000. The molecular weight of SQN2 is about 1K. The molecular weight of PMAOPEG is about 60 KDa to 200 KDa. If we take the average molecular weight of PMAOPEG, it would be around 100KDa. The Poisson distribution predicts the similar results as the observed. At mass ratio around 1: 1000, single SQN2 molecule is evolved in single PMAOPEG micelle. For the case of mass ratio at 1:20, the molar ratio is about 5:1, each PMAOPEG micelle embeds 5 SQN2 molecules.

### PA Setup

An acoustic-resolution photoacoustic microscopy (AR-PAM) system was used for this work has been reported elsewhere. Briefly, a tunable nanosecond-pulsed OPO laser (SpitLight EVO S OPO-100, InnoLas, Munich, Germany) was used to excite the samples at a repetition rate of 100 Hz. The laser beam was tuned by the half-wave plate and Glan prism and collimated by the plano-concave lens (LC4252, Thorlabs, New Jersey, USA) and plano-convex lens (LA1509, Thorlabs, New Jersey, USA), and then reshaped by the iris (D20S, Thorlabs, New Jersey, USA). After split by the PBS, the laser beam was coupled into 2 multi-mode fibers by 2 plano-convex lenses (LA1608, Thorlabs, New Jersey, USA) and 2 home-made fiber couplers, respectively. The output of the fibers illuminated the sample, and further generate PA signals. The PA signals were detected by a high-frequency focused ultrasound transducer (V214-SU, Olympus, Japan, 25 MHz central frequency). We amplified the photoacoustic signal via an US pulser-receiver (5073PR, Olympus, Japan) at 39 dB, and then the photoacoustic signal was digitized by a 2-channel data acquisition (DAQ) card (CSE1422; Gage Applied Technologies Inc., Lockport, New York) at a sampling rate of 100 MS/s. A 3-axis step motor translation stage (PSA2000-11, Zolix, Beijing, China) controlled by LabVIEW system (2011, National Instruments, TX, USA) was employed to drive the imaging head to mechanically scan over the target region, and eventually formed a 2D or 3D image.

## Acknowledgements

This work was supported by the National Natural Science Foundation of China (No. 22077135, No. 21905296).

## Author contributions

Qinchao Sun conceived and designed the experiments. Zong Chang, Liangjian Liu, Qinchao Sun performed the experiments. Qinchao Sun, Xiaojiang Xie, Chengbo Liu, Zong Chang, Liangjian Liu discussed the results. Qinchao Sun and Zong Chang analyzed the data and wrote the manuscript. All authors discussed the results and commented on the manuscript.

## Additional information

Supplementary information is available in the online version of the paper. Correspondence and requests for materials should be addressed to.

---

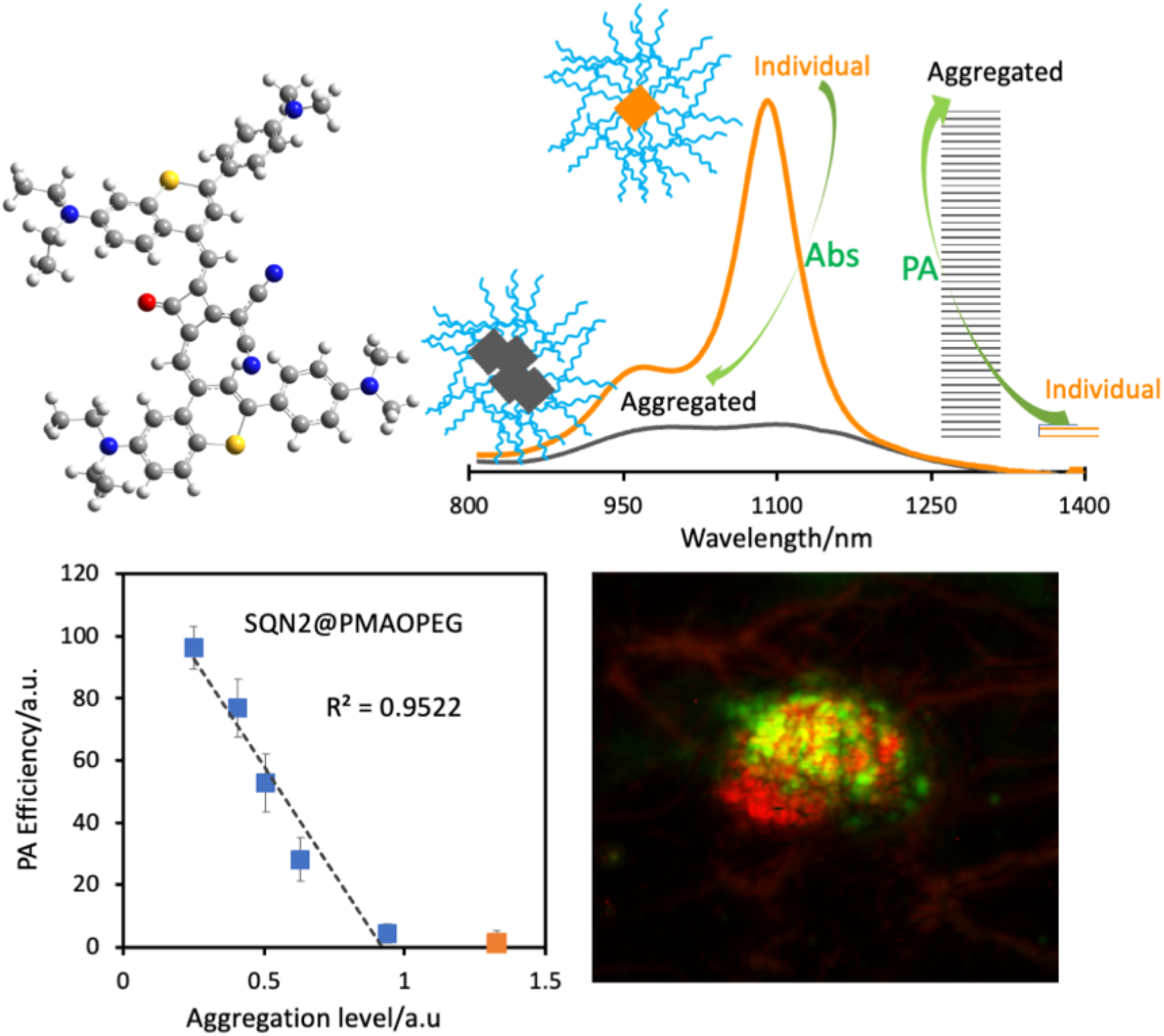

