## supplementary information for "Molecular Aggregation Induced Photoacoustics for NIR-II in vivo Imaging"

### Experimental Section

#### Instruments

$^1$H spectra were recorded on Bruker spectrometers operating at 400 MHz for 1H. For $^1$H NMR spectra, chemical shifts (δ) are reported in parts per million (ppm) with reference to tetramethylsilane (δ = 0.00 ppm) or residual protonated solvent (δ = 7.26 ppm for $CHCl_3$, δ = 3.31 ppm for $CHD_2OD$) as internal standards. Splitting patterns for $^1$H signals are designated as s (singlet), d (doublet), t (triplet), q (quartet), m (multiplet) and coupling constants (J) are reported in Hz. The UV-Vis-NIR absorption spectra were recorded on UV-2700 or UV-3600 Spectrophotometer (SHIMADZU, Japan). Fourier-transform infrared (IR) spectra were recorded for neat solid samples on a VERTEX 80 FT-IR Spectrometer (Bruker, Germany), maximum absorbances (ν) are given in $cm^{-1}$. High-resolution mass spectrometry (HRMS) data were recorded using matrix-assisted laser desorption ionization time-of-flight mass spectrometer (MALDI-TOF MS) (Bruker New ultrafleXtreme) in positive mode.

#### DFT calculation

Density functional theory (DFT) has been applied to the optimization of the ground state of all molecules in the singlet state. The calculation was performed using Becke's three-parameter exchange functional[1] combined with the Lee−Yang−Parr correlation functional[2] (B3LYP) at the level of 6-311G. The frequency calculations performed on the optimized geometries showed that they all correspond to real minima (no imaginary frequencies). All calculations were performed in

Gaussian16 program.[3]

**Cell lines**

Human cervical cancer cell line Hela cells, murine macrophage cell line RAW 264.7 cells and human prostate cancer cell line DU145 cells were provided by Research Center for Biomedical Optics and Molecular Imaging in the Shenzhen Institute of Advanced Technology, Chinese Academy of Science.

**Animal handling**

All animal handling and experimental procedures were performed in compliance with the Animal Study Committee of the Shenzhen Institutes of Advanced Technology, Chinese Academy of Sciences. All BALB/c mice were purchased from Beijing Vital River Laboratory Animal Technology Co. Ltd. During all experiments, mice were kept under anesthesia using 1.5% isoflurane mixed with pure oxygen and the body temperature was maintained at 37.5 °C by using a heating pad.

**Preparation of ZC825@BSA NPs**

100 $\mu$L ZC825 in DMSO solution (10 mg/mL) was added into BSA 1x PBS solution 4 mL, drop by drop under vortexing (the mass ratio between ZC825 and BSA was adjusted at 1: 200, 1: 100, 1: 88, 1: 50, 1: 20, respectively). The resulting solution was slowly shaken overnight at room temperature. In the end, the DMSO was removed by centrifugal filtration (Amicon 10 kDa) at speed 7000 rpm. The residue was washed three times by 1x PBS. The final solution was adjusted to 5 mL then passing through 0.22 $\mu$m syringe filter for further use.

**Preparation of SQN2@PMAOPEG**

0.5 mg SQN2 and an appropriate amount of PMAOPEG (the mass ration between SQN2 and PMAOPEG was adjusted at 1: 1000, 1: 500, 1: 200, 1: 100, 1: 20, respectively) were dissolved in $CH_2Cl_2$ in a round-bottom flask and then sonicated for 5 min. The solvent was first removed by a rotary evaporator, and then dried under vacuum from an oil pump for 3 hours to completely remove $CH_2Cl_2$. After that deionized water (2 ml) and 0.1 M NaOH (2 ml) were added. A clear solution was obtained by ultrasonication for 30 mins. The NaOH was removed by centrifugal filtration (Amicon 10 kDa). The residue was washed three times by deionized water and then five times by 1x PBS. The residue was adjusted to 5 mL by 1x PBS then passing through 0.22 $\mu$m syringe filter for further use.

Cellular Culture

The human prostate cancer cell line (DU145 cells) was cultured in RPMI 1640 medium, while the human cervical cancer cell line (Hela cells) and murine macrophage cell line (RAW 264.7 cells) were maintained in the DMEM medium. Both media were supplemented with 10% FBS and 1% (v/v) penicillin–streptomycin. The culture environment was 37 °C, and the humidification condition was 5% $CO_2$.

PA measurements of Samples

To investigate PA intensity of the ZC825@BSA and SQN2@PMAOPEG in aqueous solution at different mass ratio, a transparent plastic tube (diameter 2 mm) was applied as sample vessel. The tube was fixed to make sure the sample position unchanged during each measurement. Then ZC825@BSA aqueous solution was loaded to the tube one by one and measured with excitation at 825 nm, 3mJ/pulse. A similar procedure was applied for SQN2@PMAOPEG with excitation at 1064 nm, 3.4 mJ/pulse. For same mass concentration PA measurement with ZC825 and SQN2 25 $\mu$g/mL. For PA efficiency measurement the absorbance was adjusted to 1 with optical length 1 cm, for ZC825 at 825 nm and SQN2 at 1064 nm. The PA intensity was calculated by the integration of the maximum signal projection on Z direction.

In vitro cytotoxicity

Hela cells were dispersed within 96-well plates (1 x $10^4$ cells per well) and incubated for overnight. The cells were then exposed to ZC825@SBA or SQN2@PMAOPEG (mass ratio at 1: 20) at various concentrations (0-200 $\mu$g/mL referring to ZC825 or SQN2) for 12 hours. The cell viabilities were measured using the standard CCK-8 assay, as instructed by the manufacturer.

In vivo cytotoxicity

ZC825@BSA or SQN2@PMAOPEG (mass ratio 1:20) at a dose of 2 mg/kg or saline were intravenously administered into tail vein of healthy BALB/c mice (3 mice per group), respectively. After fourteen days, the hearts, livers, kidneys, lungs, and spleen were harvested. After fixation in 10% neutral buffered formalin, embedding into paraffin and sectioning at 5 µm thickness, the tissues were stained with H&E and examined by means of a digital microscope (Olympus, CX31, Japan).

Mouse tumor model

8 weeks old BALB/c nude mice were used for tumor implantation. DU145 cells were incubated with RPMI 1640 medium (10% FBS and 1% penicillin-streptomycin) and ZC825@BSA (ZC825

concentration 250 µg/mL, mass ratio 1: 20) for 2 hours. The stained DU145 cells (about 1 x $10^6$) were suspended in 100 µL 1x PBS and then inoculated subcutaneously into the posterior flank of mouse right hindlimb under full anesthesia. After about one week, the inoculated stained cells grew up to lump with diameter about 3 mm.

In vivo PA imaging

Macrophage cells (RAW264.7) were cultured in DMEM medium with 10% FBS and 1% penicillin-streptomycin in the presence of SQN2@PMAOPEG aqueous solution (SQN2 concentration 250 µg/mL, mass ratio 1: 20) for 2 hours. The stained macrophage cells were collected and suspended in 1x PBS. Hela cells were cultured in DMEM medium with 10% FBS and 1% penicillin-streptomycin in the presence of ZC825@BSA (ZC825 concentration 250 µg/mL, mass ratio 1: 20) for 2 hours. After that the stained Hela cells were collected and suspended in 1x PBS.

(1) The above stained macrophage cells and Hela cells were mixed with matrix gel, and then subcutaneously injected into the right hindlimb at different location. The PA images were recorded with excitation at 825 nm (3 mJ/pulse) and 1064 nm (15 mJ/pulse).

(2) For tumor model imaging, the above stained macrophages cells in 1x PBS were subcutaneously injected into the location of the tumor. The PA images were recorded with excitation at 825 nm (3 mJ/pulse) and 1064 nm (15 mJ/pulse).

(3) For blood and lymph vessels imaging, 10 µL SQN2@PMAOPEG (SQN2 concentration 1.0 mg/mL, mass ratio 1:20) was subcutaneously injected into the left hind paw. After 30 mins, 200 µL ZC825@BSA (ZC825 concentration 1.0 mg/mL, mass ratio 1:20) was intravenously injected into the tail vein. The PA imaging was conducted sooner after injection of ZC825@BSA with excitation at 825 nm (3 mJ/pulse). After that the PA image was conducted with excitation at 1064 nm (15 mJ/pulse).

General synthetic route of dyes

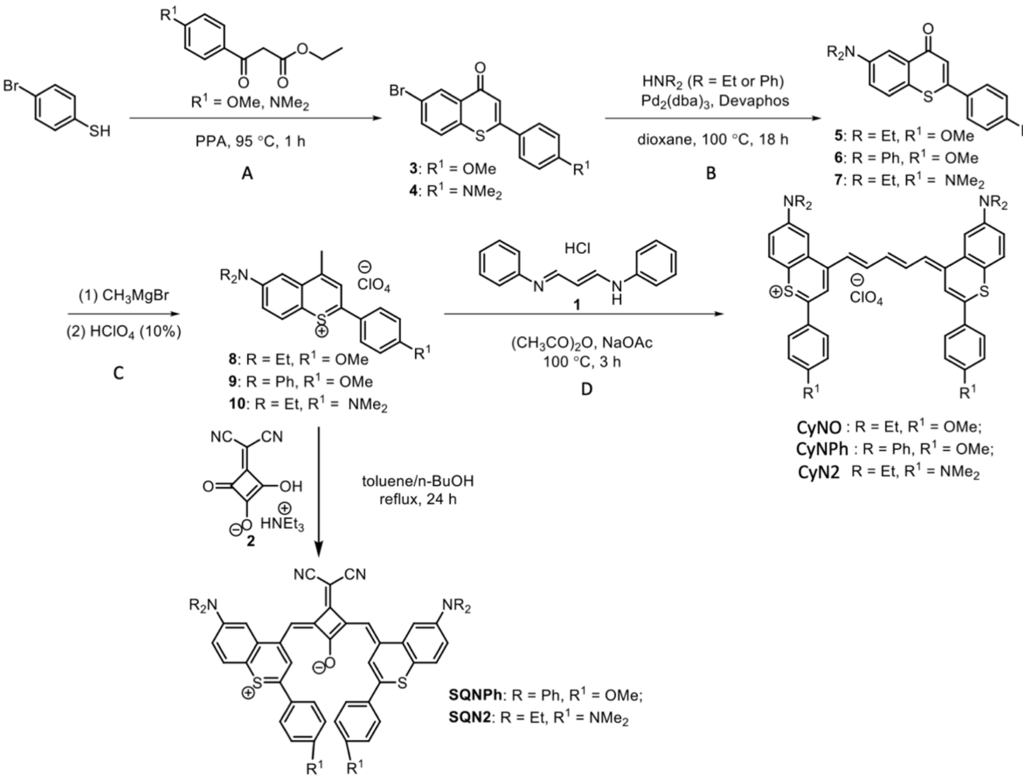


**General procedure A:** The synthetic procedure was according to reported literature with a bit modification.[4] To a hot polyphosphoric acid (PPA), ethyl 3-oxo-3-phenylpropanoate and 3-bromothiophenol were added gradually, the mixture was stirred vigorously at 95 °C for 1 h. After cooling to room temperature, ice water was added slowly to quench the reaction and extracted with $CH_2Cl_2$ (3 times). The combined organic extracts were dried on $Na_2SO_4$, the solvent was evaporated. The crude product was further purified by a flash column chromatography on silica gel to give the desired product.

**General procedure B:** To a Schlenk flask, compound 3 or 4, $Pd_2(dba)_3$, DavePhos, and $Cs_2CO_3$ were added gradually under argon atmosphere. Anhydrous dioxane was added, following the addition of diethylamine or diphenylamine. The reaction was stirred at 100 °C for 18 h. After cooling to room temperature, the mixture was filtered over celite, and the filtrate was evaporated. The crude product was further purified by a column chromatography on silica gel to give the product.

**General procedure C:** To a solution of compound 5 (6 or 7) in anhydrous THF, 1.0 M $CH_3MgBr$ was added dropwise and allowed to stir at room temperature for 2 h. The solution was poured into 10% aqueous $HClO_4$ and extracted with $CH_2Cl_2$ (3 times). The combined organic extracts were dried over

$Na_2SO_4$, filtered, and evaporated to give the desired product without further purification.

**General procedure D:** To a mixture of compound 8 (9 or 10), N-((1E,3E)-3-(phenylimino)prop-1-en-1-yl)aniline hydrochloride and NaOAc, MeCN and $Ac_2O$ were added in a sequence. The mixture was heated at 100 °C under nitrogen for 3 h. After cooling to room temperature, the mixture was evaporated and purified by a column chromatography on silica gel to give the final product.

### Synthesis of CyNO

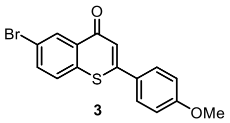

**3**

6-bromo-2-(4-methoxyphenyl)-4H-thiochromen-4-one 3: Followed the general procedure A, using 4-bromobenzenethiol (2.0 g, 10.6 mmol) with ethyl 3-(4-methoxyphenyl)-3-oxopropanoate (2.6 g, 11.7 mmol) and PPA (10 mL). flash chromatography (Petroleum ether/$CH_2Cl_2$, 50/50 to 0/100) afforded the compound 3 (1.18 g, 32%). $^1H$ NMR (400 MHz, $CDCl_3$) δ 8.66 (d, J = 2.0 Hz, 1H), 7.70 (dd, J = 2.0, 8.5 Hz, 1H), 7.63 (d, J = 8.8 Hz, 2H), 7.51 (d, J = 8.5 Hz, 1H), 7.19 (s, 1 H), 7.00 (d, J = 8.8 Hz, 2H), 3.87 (s, 3H).

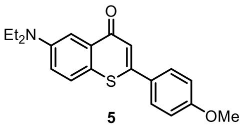

**5**

6-(diethylamino)-2-(4-methoxyphenyl)-4H-thiochromen-4-one 5: Followed the general procedure B, using compound 3 (174 mg, 0.5 mmol), $HNEt_2$ (0.26 mL, 2.5 mmol), $Pd_2(dba)_3$ (11.4 mg, 0.013 mmol), Davephos (4.9 mg, 0.013 mmol), $Cs_2CO_3$ (407 mg, 1.25 mmol) and anhydrous dioxane (2.5 mL). flash chromatography (Petroleum ether/$CH_2Cl_2$, 50/50; then EtOAc/$CH_2Cl_2$, 4/96) afforded the compound 5 (100 mg, 60%) as a yellow solid. $^1H$ NMR (400 MHz, $CDCl_3$) δ 7.72 (d, J = 2.9 Hz, 1H), 7.64 (d, J = 8.8 Hz, 2H), 7.46 (d, J = 8.9 Hz, 1H), 7.16 (s, 1 H), 7.22 (d, J = 9.0, 2.9 Hz, 1H), 6.97 (d, J = 8.8 Hz, 2H), 3.85 (s, 3H), 3.45 (q, J = 7.0 Hz, 4H), 1.20 (t, J = 7.0 Hz, 6H).

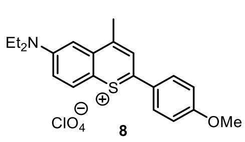


6-(diethylamino)-2-(4-methoxyphenyl)-4-methylthiochromenylium perchlorate 8: Followed the general procedure C, using compound 5 (68 mg, 0.2 mmol), 1.0 M $CH_3MgBr$ (0.6 mL, 0.6 mmol) and anhydrous THF (2.0 mL). Compound 8 (83 mg, 95%) was obtained as a green solid. $^1$H NMR (400 MHz, $CDCl_3$) δ 8.43 (s, 1H), 8.22 (d, J = 9.2 Hz, 1H), 7.99 (d, J = 8.2 Hz, 2H), 7.49 (d, J = 8.8 Hz, 1H), 7.10 (s, 1H), 6.99 (d, J = 8.2 Hz, 2H), 3.78 (s, 3H), 3.55 (q, J = 6.8 Hz, 4H), 2.99 (s, 3H), 1.27 (t, J = 6.8 Hz, 6H).

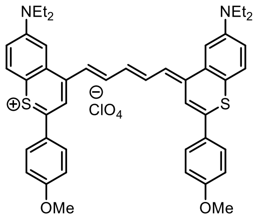


6-(diethylamino)-4-((1E,3E)-5-((E)-6-(diethylamino)-2-(4-methoxyphenyl)-4H-thiochromen-4-ylidene)penta-1,3-dien-1-yl)-2-(4-methoxyphenyl)thiochromenylium perchlorate CyNO: Followed the general procedure D, using compound 8 (66 mg, 0.15 mmol), N-((1E,3E)-3-(phenylimino)prop-1-en-1-yl)aniline hydrochloride (19.5 mg, 0.075 mmol), NaOAc (12.3 mg, 0.15 mmol), anhydrous $Ac_2O$ (0.3 mL), anhydrous acetonitrile (0.6 mL). Flash chromatography (MeOH/$CH_2Cl_2$, 0/100 to 1/99) afforded CysNO (37 mg, 30%). $^1$H NMR (400 MHz, $CDCl_3$) δ 8.20 (t, J = 12.9 Hz, 2H), 8.03 (s, 2H), 7.77 (d, J = 8.6 Hz, 4H), 7.35-7.16 (m, 5H), 7.14 (s, 2H), 6.98 (d, J = 8.6 Hz, 4H), 6.77 (m, 2H), 3.86 (s, 6H), 3.32 (q, J = 6.8 Hz, 8H), 1.17 (t, J = 6.8 Hz, 12H). HRMS (MALDI-TOF) m/z: $[M\text{-}ClO4]^+$ calc. for $C_{45}H_{47}N_2O_2S_2^+$: 711.307; Found: 711.163.

### Synthesis of CyNPh

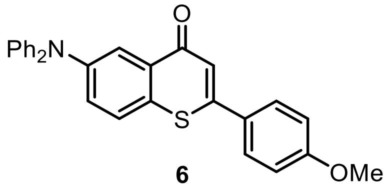


6-(diphenylamino)-2-(4-methoxyphenyl)-4H-thiochromen-4-one 6: Followed the general procedure

B, using compound 3 (174 mg, 0.5 mmol), $HNPh_2$ (423 mg, 2.5 mmol), $Pd_2(dba)_3$ (11.4 mg, 0.013 mmol), Davephos (4.9 mg, 0.013 mmol), $Cs_2CO_3$ (407 mg, 1.25 mmol) and anhydrous dioxane (2.5 mL). flash chromatography (Petroleum ether/EtOAc, 80/20) afforded the compound 6 (151 mg, 70%) as a yellow solid. $^1H$ NMR (400 MHz, $CDCl_3$) δ 8.19 (d, J = 2.5 Hz, 1H), 7.64 (d, J = 8.8 Hz, 2H), 7.47 (d, J = 8.7 Hz, 1H), 7.35 (dd, J = 8.7, 2.5 Hz, 1H), 7.31-7.24 (m, 4H), 7.15-7.10 (m, 5H), 7.10-7.04 (m, 2H), 7.02-6.97 (m, 2H), 3.87 (s, 3H).

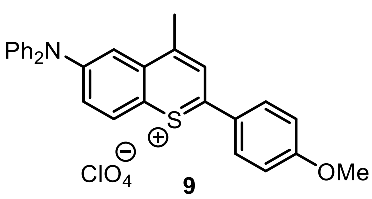

**9**

6-(diphenylamino)-2-(4-methoxyphenyl)-4-methylthiochromenylium perchlorate 9: Followed the general procedure C, using compound 6 (87 mg, 0.2 mmol), 1.0 M $CH_3MgBr$ (0.6 mL, 0.6 mmol) and anhydrous THF (2.0 mL). Compound 9 (103 mg, 96%) was obtained as a black solid. $^1H$ NMR (400 MHz, $CDCl_3$) δ 8.53 (s, 1H), 8.21 (d, J = 9.5 Hz, 1H), 8.08 (d, J = 8.9 Hz, 2H), 7.67-7.60 (m, 2H), 7.44 (t, J = 7.8 Hz, 4H), 7.32-7.24 (m, 6H), 6.99 (d, J = 8.9 Hz, 2H), 3.84 (s, 3H), 2.79 (s, 3H).

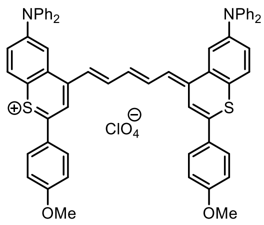


6-(diphenylamino)-4-((1E,3E)-5-((E)-6-(diethylamino)-2-(4-methoxyphenyl)-4H-thiochromen-4-ylidene)penta-1,3-dien-1-yl)-2-(4-methoxyphenyl)thiochromenylium perchlorate CyNPh: Followed the general procedure D, using compound 9 (80 mg, 0.15 mmol), N-((1E,3E)-3-(phenylimino)prop-1-en-1-yl)aniline hydrochloride (19.5 mg, 0.075 mmol), NaOAc (12.3 mg, 0.15 mmol), anhydrous $Ac_2O$ (0.3 mL), anhydrous acetonitrile (0.6 mL). Flash chromatography ($MeOH/CH_2Cl_2$, 0/100 to 1/99) afforded CysNPh (41 mg, 27%). $^1H$ NMR (400 MHz, $CDCl_3$) δ 8.53 (t, J = 12.9 Hz, 2H), 8.30 (s, 2H), 7.97 (d, J = 8.8 Hz, 4H), 7.71 (s, 2H), 7.50 (d, J = 8.8 Hz, 4H), 7.36 (t, J = 7.8 Hz, 8H), 7.19-7.12 (m, 12H), 7.05 (d, J = 8.8 Hz, 4H), 6.59 (d, J = 13.3 Hz, 2H), 6.44 (t, J = 12.5 Hz, 1H), 3.84 (s, 6H). HRMS (MALDI-TOF) m/z: [M-ClO4]$^+$ calc. for $C_{61}H_{47}N_2O_2S_2^+$: 903.307; Found: 903.296.

Synthesis of CyN2

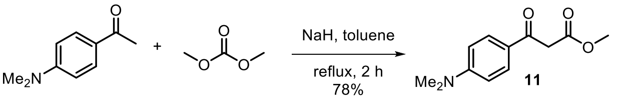

methyl 3-(4-(dimethylamino)phenyl)-3-oxopropanoate 11: The procedure of the synthesis of 11 was according to reported literature with a bit modification.[5] To a suspension of NaH (60% dispersion in mineral oil, 1.0 g, 25.2 mmol) in toluene (10 mL), dimethyl carbonate (1.52 mL, 1.62 g, 18.0 mmol) was added and heated to reflux. 1-(4-(dimethylamino)phenyl)ethan-1-one (1.47 g, 9.0 mmol) in toluene (10 mL) was added dropwise during 30 min and the mixture was refluxed for 4 h. After cooling to room temperature, the reaction mixture was poured onto ice water (25 mL) and aq. HCl solution (1.0 M, 25 mL) and extracted with AcOEt (3 × 25 mL). The combined organic layers were washed with brine (25 mL), dried over anhydrous $MgSO_4$ and concentrated under reduced pressure. The residue was purified by a flash chromatography (EtOAc/petroleum ether, 50:50) to afford the compound 11 (1.56 g, 78%) as a yellow solid. $^1$H NMR (400 MHz, $CDCl_3$) δ 7.86 (d, J = 8.2 Hz, 2H), 6.66 (d, J = 8.2 Hz, 2H), 3.94 (s, 2H), 3.75 (s, 3H), 3.08 (s, 6H).

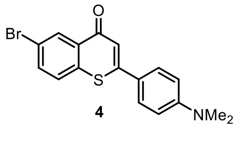

6-bromo-2-(4-(dimethylamino)phenyl)-4H-thiochromen-4-one 4: Followed the general procedure A, using 4-bromobenzenethiol (567 mg, 3 mmol) with compound 11 (729 mg, 3.3 mmol) and PPA (6.0 mL). flash chromatography ($CH_2Cl_2$, 100; Petroleum ether/EtOAc, 75/25 to 50/50) afforded the compound 4 (303 mg, 28%). $^1$H NMR (400 MHz, $CDCl_3$) δ 8.62 (d, J = 1.9 Hz, 1H), 7.64 (dd, J = 8.5, 1.9 Hz, 1H), 7.56 (d, J = 8.8 Hz, 2H), 7.46 (d, J = 8.5 Hz, 1H), 7.17 (s, 1H), 6.69 (d, J = 8.9 Hz, 2H), 3.03 (s, 6H).

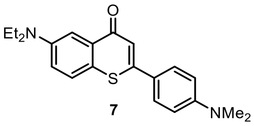


**7**

6-(diethylamino)-2-(4-(dimethylamino)phenyl)-4H-thiochromen-4-one 7: Followed the general procedure B, using compound 4 (1.08 g, 3.0 mmol), $HNEt_2$ (1.54 mL, 15 mmol), $Pd_2(dba)_3$ (137 mg, 0.15 mmol), Davephos (60 mg, 0.15 mmol), $Cs_2CO_3$ (2.44 g, 7.5 mmol) and anhydrous dioxane (15 mL). flash chromatography (Ether/$CH_2Cl_2$, 0/100 to 2/98) afforded the compound 7 (767 mg, 73%) as a yellow solid. $^1$H NMR (400 MHz, $CDCl_3$) δ 7.75 (d, J = 2.8 Hz, 1H), 7.63 (d, J = 8.8 Hz, 2H), 7.48 (d, J = 8.9 Hz, 1H), 7.20 (s, 1H), 7.03 (dd, J = 8.9, 2.8 Hz, 1H), 6.74 (d, J = 8.8 Hz, 2H), 3.47 (q, J = 7.0 Hz, 4H), 3.03 (s, 6H), 1.22 (t, J = 7.0 Hz, 6H).

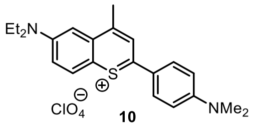


**10**

6-(diethylamino)-2-(4-(dimethylamino)phenyl)-4-methylthiochromenylium perchlorate 10: Followed the general procedure C, using compound 7 (408 mg, 1.16 mmol), 1.0 M $CH_3MgBr$ (3.48 mL, 3.48 mmol) and anhydrous THF (30 mL). Compound 10 (517 mg, 99%) was obtained as a black solid. $^1$H NMR (400 MHz, $CDCl_3$) δ 8.31 (s, 1H), 7.98 (dd, J = 14.1, 9.2 Hz, 3H), 7.31 (dd, J = 9.2, 2.6 Hz, 1H), 7.06 (d, J = 2.6 Hz, 1H), 6.78 (d, J = 9.2 Hz, 2H), 3.52 (q, J = 7.1 Hz, 4H), 3.15 (s, 6H), 2.91 (s, 3H), 1.27 (t, J = 7.1 Hz, 6H).

### Synthesis of CyN2

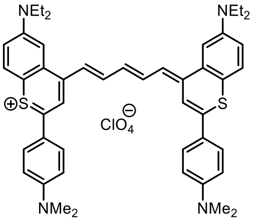


6-(diethylamino)-4-((1E,3E)-5-((E)-6-(diethylamino)-2-(4-(dimethylamino)phenyl)-4H-thiochromen-4-ylidene)penta-1,3-dien-1-yl)-2-(4-(dimethylamino)phenyl)thiochromenylium CyN2: Followed the general procedure D, using compound 10 (45 mg, 0.10 mmol), N-((1E,3E)-3-(phenylimino)prop-1-en-1-yl)aniline hydrochloride (13 mg, 0.05 mmol), NaOAc (22 mg, 0.27 mmol), anhydrous $Ac_2O$ (1.0 mL). Flash chromatography (MeOH/$CH_2Cl_2$, 0/100 to 2/98) afforded CysN2 (20 mg, 27%). $^1H$ NMR (400 MHz, MeOD) δ 8.2-7.95 (m, 4H), 7.93-7.84 (m, 5H), 7.82-7.74 (m, 3H), 7.70-7.50 (m, 2H), 7.25-6.95 (m, 5H), 6.88 (t, J = 12.3 Hz, 2H), 3.80-3.55 (m, 8H), 3.08 (s, 12H), 1.26 (t, J = 6.9 Hz, 12H). ($^1H$ NMR data is the proton spectra of CysN2 hydrochloride). HRMS (MALDI-TOF) m/z: $[M-ClO4]^+$ calc. for $C_{47}H_{53}N_4S_2^+$: 737.371; Found: 737.326

### Synthesis of SQNPh

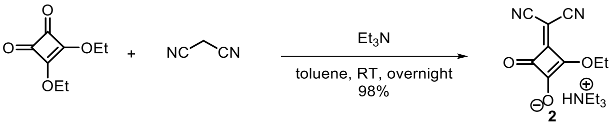


triethyl-λ[4]-azane, 3-(dicyanomethylene)-2-hydroxy-4-oxocyclobut-1-en-1-olate salt 2: The synthesis of 2 was based on a literature procedure.[6] To a solution of malononitrile (165 mg, 2.5 mmol) in toluene (10 mL), 3,4-diethoxycyclobut-3-ene-1,2-dione (0.38 mL, 2.55 mmol) was added dropwise. $Et_3N$ (0.35 mL, 2.55 mmol) was added slowly at room temperature. The reaction was allowed to stir for overnight, the resulting crystalline was filtrated and washed with toluene and ether to give the compound 2 (650 mg, 99%). $^1H$ NMR (400 MHz, MeOD) δ 4.70 (q, J = 7.1 Hz, 2H), 3.22 (q, J = 7.3 Hz, 6H), 1.44 (t, J = 7.1 Hz, 3H), 1.31 (t, J = 7.3 Hz, 6H).

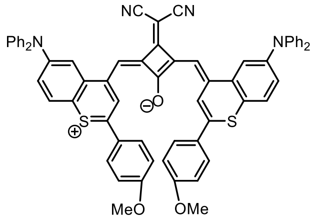


**SQNPh**

(Z)-3-(dicyanomethylene)-2-(((E)-6-(diphenylamino)-2-(4-methoxyphenyl)-4H-thiochromen-4-ylidene)methyl)-4-((6-(diphenylamino)-2-(4-methoxyphenyl)thiochromenylium-4-yl)methylene)cyclobut-1-en-1-olate SQNPh: To a solution of compound 9 (107 mg, 0.20 mmol) in toluene/n-BuOH (v/v = 3/1, 10 mL), compound 2 (29 mg, 0.10 mmol) was added in one portion. The reaction was stirred for 24 h under reflux. The solvents were evaporated and the residue was purified by column chromatography on silica gel (MeOH/$CH_2Cl_2$, 0/100 to 10/90) to provide SQNPh

(61 mg, 31%) as a black solid. $^{1}H$ NMR (400 MHz, $CDCl_3$) δ 9.49 (s, 2H), 7.99 (d, J = 8.4 Hz, 4H), 7.82 (s, 2H), 7.42-7.30 (m, 12H), 7.15 (d, J = 7.6 Hz, 12H), 7.04 (s, 2H), 6.94 (d, J = 8.3 Hz, 4H), 3.84 (s, 6H). HRMS (MALDI-TOF) m/z: $[M+H]^+$ calc. for $C_{65}H_{45}N_4O_3S_2^+$: 993.293; Found: 993.180.

### Synthesis of SQN2

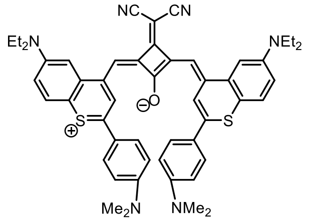


SQN2

(Z)-3-(dicyanomethylene)-2-(((E)-6-(diethylamino)-2-(4-(dimethylamino)phenyl)-4H-thiochromen-4-ylidene)methyl)-4-((6-(diethylamino)-2-(4-(dimethylamino)phenyl)thiochromenylium-4-yl)methylene)cyclobut-1-en-1-olate SQN2: To a solution of compound 10 (90 mg, 0.20 mmol) in toluene/n-BuOH (v/v = 3/1, 10 mL), compound 2 (29 mg, 0.10 mmol) was added in one portion. The reaction was stirred for 24 h under reflux. The solvents were evaporated, and the residue was purified by column chromatography on silica gel (MeOH/$CH_2Cl_2$, 0/100 to 10/90) to provide SQN2 (46 mg, 28%) as a black solid. No $^{1}H$ NMR (400 MHz, $CDCl_3$) spectrum was obtained as the low solubility in DCM. HRMS (MALDI-TOF) m/z: $[M+H]^+$ calc. for $C_{51}H_{51}N_6OS_2^+$: 827.356; Found: 827.205.

### Synthesis of ZC825

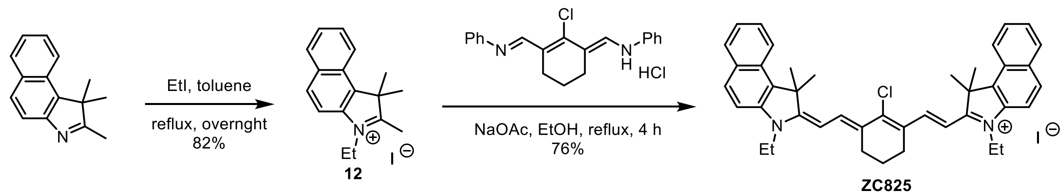


3-ethyl-1,1,2-trimethyl-1H-benzo[e]indol-3-ium iodide 12: The synthesis of 12 was Based on a literature procedure.[7] To a solution of 1,1,2-trimethyl-1H-benzo[e]indole ( 1.05 g, 5.0 mmol) in toluene (10 mL), EtI (0.48 mL, 6.0 mmol) was added under argon atmosphere. The reaction mixture was stirred under reflux for overnight, then allowed to cool to room temperature. The solvent was removed under vacuum filtration and the residue was washed with ether to afford the desired product 12 (1.5 g, 82%) as a blue solid. $^1$H NMR (400 MHz, $CDCl_3$) δ 8.16-8.03 (m, 3H), 7.81 (d, J = 8.9 Hz, 1H), 7.74 (t, J = 7.4 Hz, 1H), 7.68 (t, J = 7.4 Hz, 1H), 4.89 (q, J = 7.4 Hz, 2H), 3.24 (s, 3H), 1.88 (s,6H), 1.67 (t, J = 7.4 Hz, 3H).

2-((E)-2-((E)-2-chloro-3-((E)-2-(3-ethyl-1,1-dimethyl-1,3-dihydro-2H-benzo[e]indol-2-ylidene)ethylidene)cyclohex-1-en-1-yl)vinyl)-3-ethyl-1,1-dimethyl-1H-benzo[e]indol-3-ium iodide ZC825: To a solution of compound 12 (182.5 mg, 0.5 mmol) in EtOH (10 mL), sodium acetate (41 mg, 0.5 mmol) and N-((E)-(2-chloro-3-((E)-(phenylimino)methyl)cyclohex-2-en-1-ylidene)methyl)aniline hydrochloride (90 mg, 0.25 mmol) was added gradually under argon atmosphere. The reaction mixture was stirred under reflux for 4 h, then allowed to cool to room temperature. After evaporation of solvent, the residue was purified by flash chromatography (MeOH/acetone/$CH_2Cl_2$, 2.5/12.5/85) to afford ZC825 (140 mg, 76%) as a green solid. $^1$H NMR (400 MHz, MeOD) δ 8.59 (d, J = 14.3 Hz, 2H), 8.30 (d, J = 8.5 Hz, 2H), 8.07 (d, J = 8.8 Hz, 2H), 8.03 (d, J = 8.2 Hz, 2H), 7.71-7.63 (m, 4H), 7.56-7.50 (m, 2H), 6.37 (d, J = 14.2 Hz, 2H), 4.38 (q, J = 7.2 Hz, 4H), 2.81 (t, J = 6.1 Hz, 4H), 2.07-2.01 (m, 14H), 1.51 (t, J = 7.2 Hz, 6H). HRMS (MALDI-TOF) m/z: $[M-I]^+$ calc. for $C_{42}H_{44}ClN_2^+$: 611.312; Found: 611.271.

### Preparation of PMAOPEG

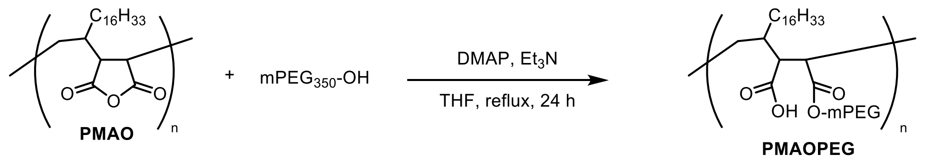


PMAOPEG: To a solution of PMAO (1.0 g, 30- 50 KDa) in anhydrous THF (100 mL), $mPEG_{350}$ (10 g), DMAP (122 mg, 2.7 mmol) and $Et_3N$ (7.8 mL, 54.6 mmol) were added gradually, the reaction was refluxed for 24 h. The solvent was evaporated, the residue was dried on oil pump for 3 hours. Water was added and the resulting solution was dialyzed with a 10 KDa dialysis bag for one month, evaporation of solvent to provide PMAOPEG (1.4 g) as a pale-yellow solid. The molecular weight of PMAOPEG should be around 60-100 KDa in the case of half carboxylate groups were conjugated with mPEG.

IR(PMAO) ν 2922, 2852, 1851, 1778, 1710, 1466, 1219, 1072, 923 ($cm^{-1}$).

IR(PMAOPEG) ν 2920, 2852, 1724, 1647, 1560, 1465, 1215, 1175, 1103, 945 ($cm^{-1}$).

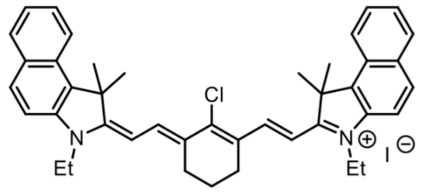


**ZC825**

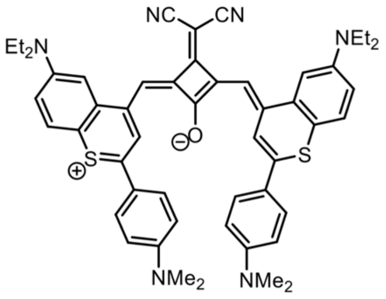


**SQN2**

Figure S1. The molecular structure of the ZC825 and SQN2.

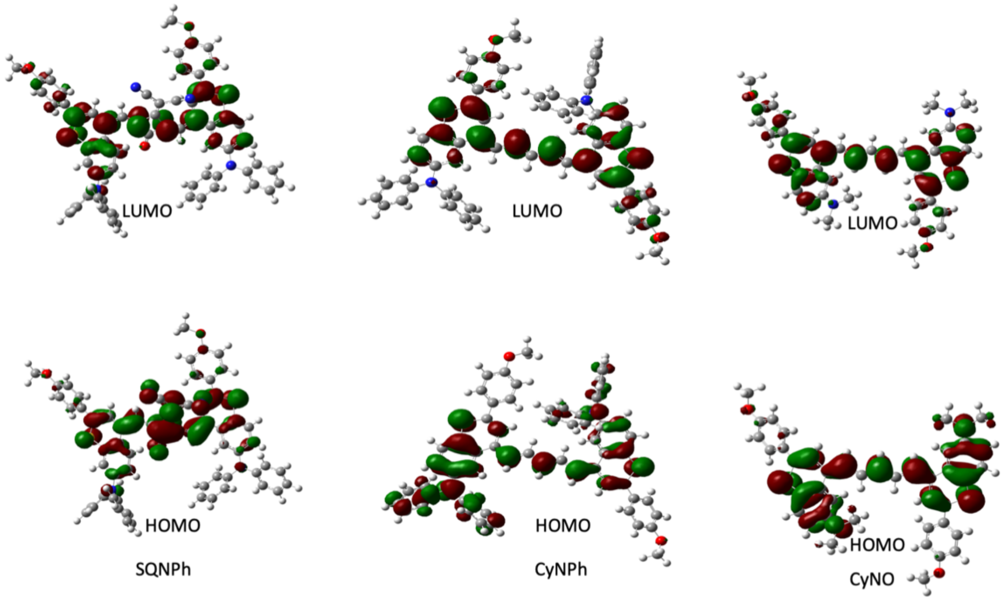


Figure S2. The visualized HOMO and LUMO molecular orbitals of the SQNPh, CyNPh and CyNO.

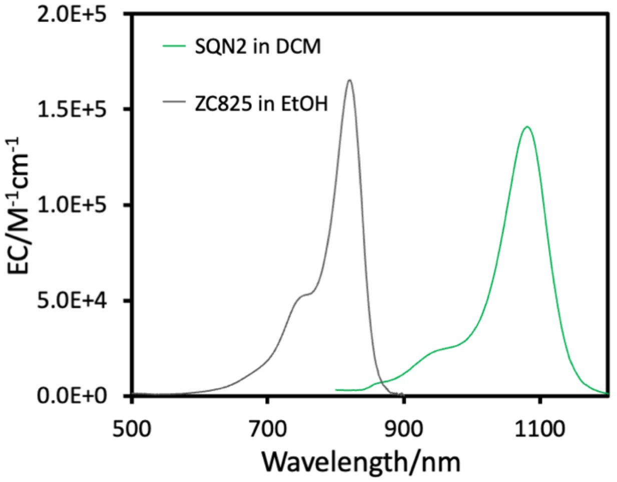


Figure S3. The molar absorption extinction coefficient (EC) of ZC825 in EtOH and SQN2 in DCM.

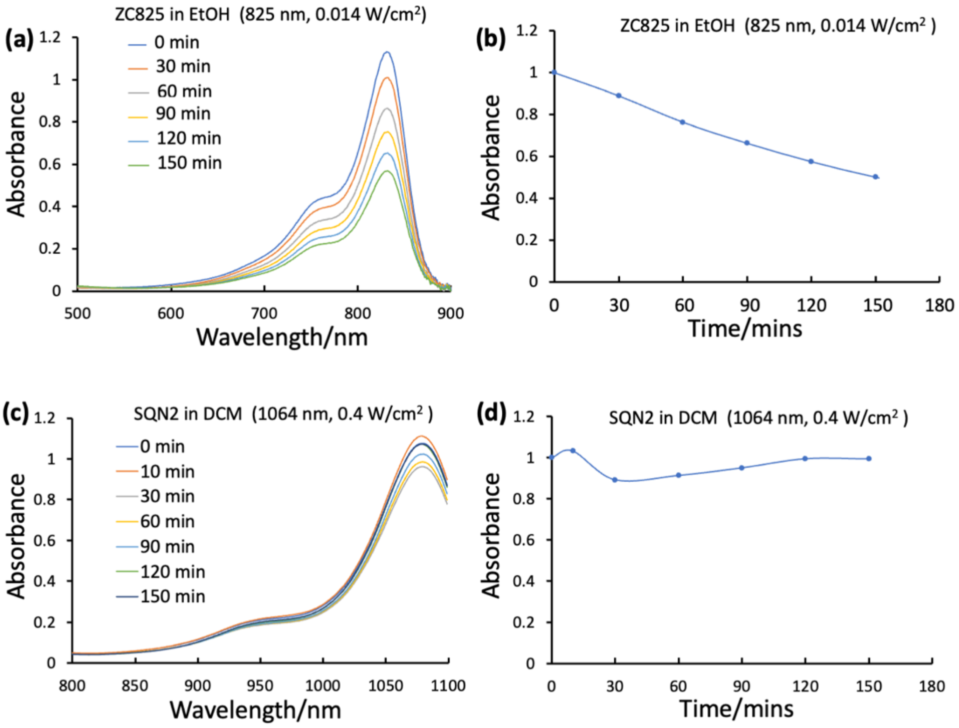


Figure S4. The photostability of ZC825 and SQN2. (a) The absorption spectra of ZC825 in EtOH at different laser irradiation time, excitation 825 nm, 0.014W/cm$^2$. (b) The absorbance variation of ZC825 at the absorption maximum as a function of laser irradiation time, excitation 825 nm, 0.014W/cm$^2$. (a) The absorption spectra of SQN2 in DCM at different laser irradiation time, excitation 1064 nm, 0.4W/cm$^2$. (b) The absorbance variation of SQN2 at the absorption maximum as a function of laser irradiation time, excitation 1064 nm, 0.4W/cm$^2$.

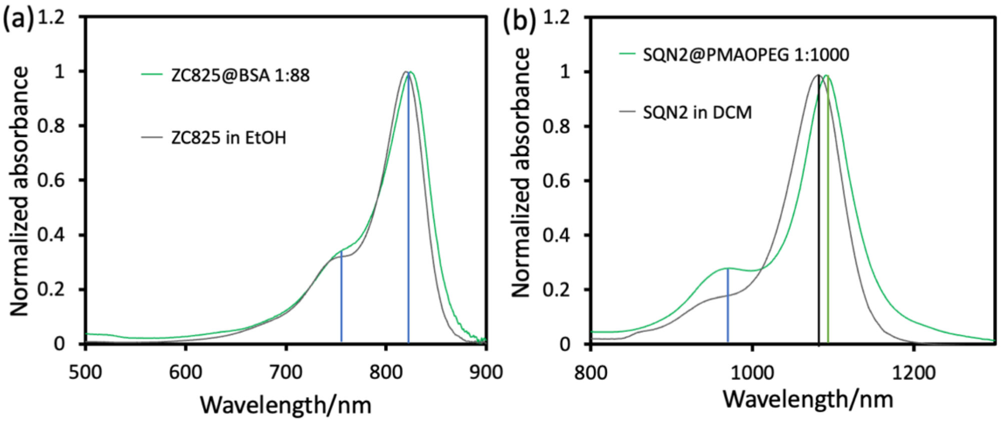


Figure S5. The normalized absorption spectra of ZC825 and SQN2. (a) The normalized absorption spectra of ZC825 in EtOH (grey curve) and the ZC825@BSA at mass ratio about 1:88 (green curve), the vertical line indicates the absorption peak and shoulder. (b) The normalized absorption spectra of SQN2 in DCM (grey curve) and the SQN2@PMAO at the mass ratio about 1:1000 (green curve), the vertical line indicates the absorption peak and shoulder.

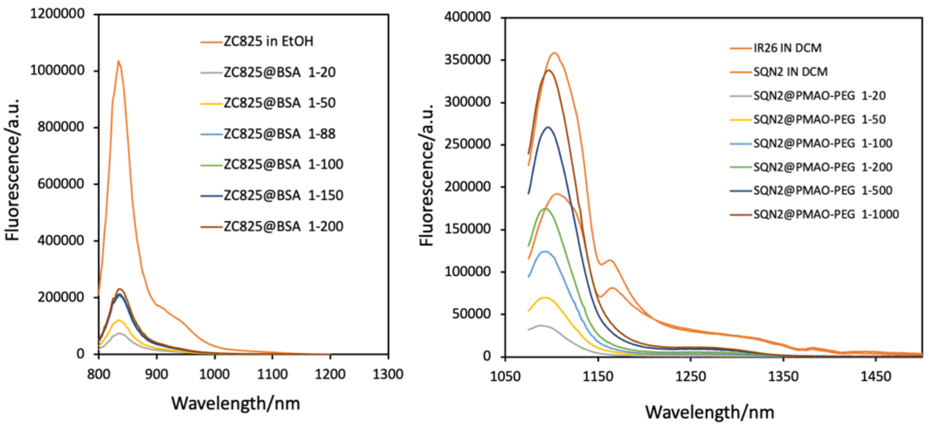


Figure S6. The fluorescence spectra of ZC825 in EtOH and ZC825@BSA at different mass ratio in aqueous solution with absorbance about 0.1 (optical path 1 cm) with excitation 825 nm and the IR26 in DCM, SQN2 in DCM, SQN2@PMAOPEG at different mass ratio in aqueous solution with absorbance about 0.1 (optical path 1 cm) with excitation at 1064 nm. The fluorescence quantum yield of ZC825 in EtOH about 4% and of IR26 in DCM about 0.03%.

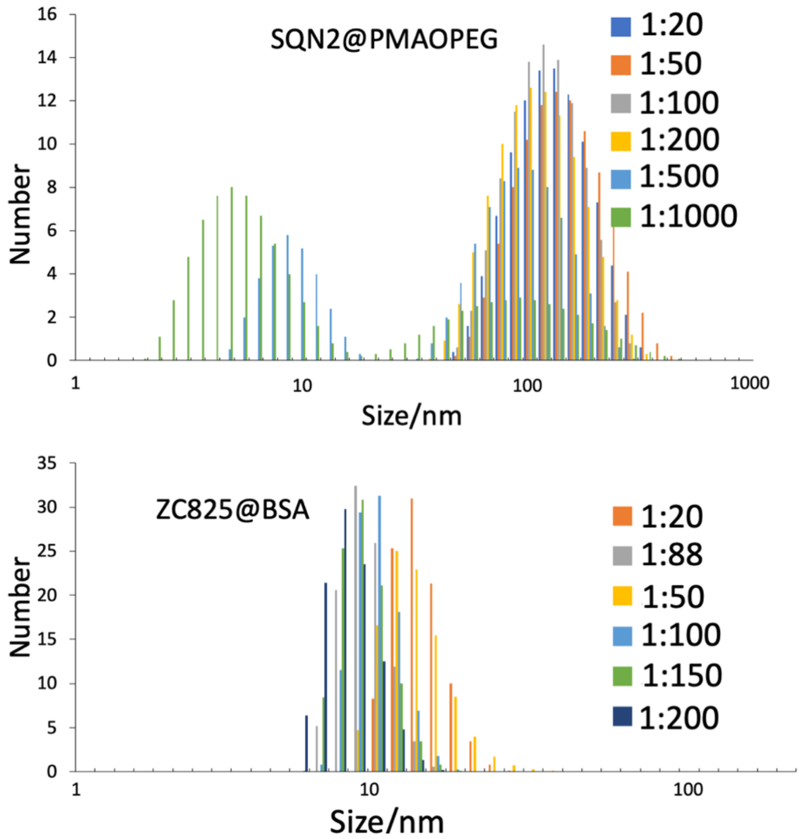

Figure S7. The hydrodynamic size distribution of SQN2@PMAOPEG and ZC825@BSA at different mass ratio. There is a distribution of hydrodynamic size around few nanometers of SQN2@PMAOPEG at mass ratio of 1:1000 and 1:500, which might come from the truth that according to Poisson distribution, the large amount of PMAOPEG without binding with SQN2 molecules.

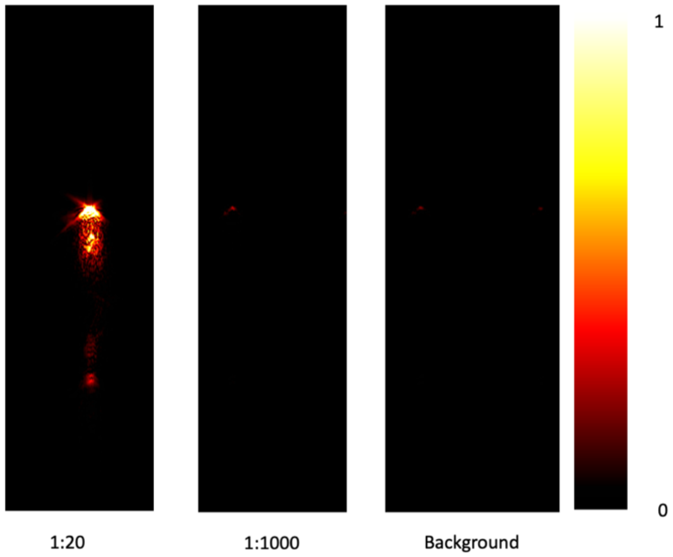


Figure S8. The PA image of SQN2@PMAOPEG with different mass ratio of SQN2 and PMAOPEG under the same PMAOPEG concentration about 200 μg/mL under excitation at 1064 nm, 15 mJ/pulse.

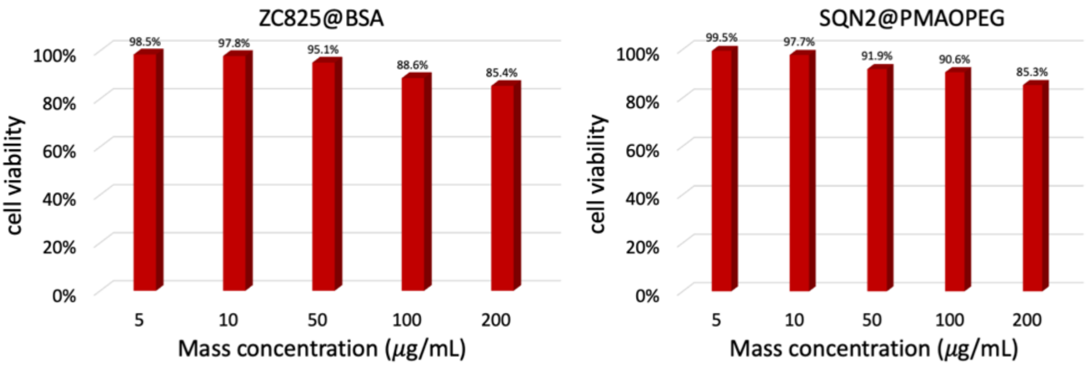


Figure S9. The cell viability measurement at different mass concentration of ZC825 and SQN2 with ZC825@BSA and SQN2@PMAOPEG at mass ratio about 1: 20.

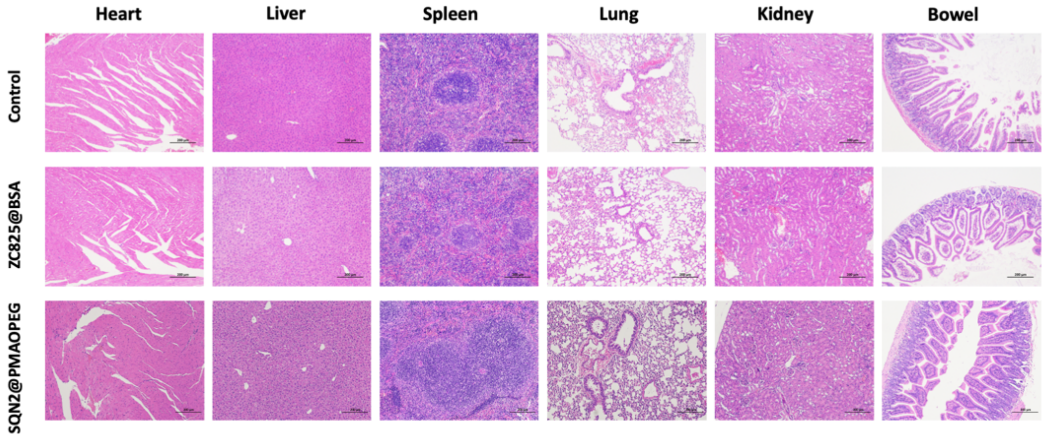


Figure S10. The representative H&E-stained sections of the heart, liver, spleen, lungs, and kidneys collected from the mice with intravenous administration of saline (control), ZC825@BSA and SQN2@PMAOPEG mass ratio about 1: 20.

References:

1. Becke, A. D. Density-functional thermochemistry. III. The role of exact exchange. J. Chem. Phys. 1993, 98, 5648−5652.
2. Lee, C., Yang, W., Parr, R. G. Development of the Colle-Salvetti correlation-energy formula into a functional of the electron density. Phys. Rev. B. 1988, 37, 785−789.
3. Gaussian 16, Revision C.01, Frisch, M. J.; Trucks, G. W.; Schlegel, H. B.; Scuseria, G. E.; Robb, M. A.; Cheeseman, J.R.; Scalmani, G.; Barone, V.; Mennucci, B.; Petersson, G. A.; Nakatsuji, H.; Caricato, M.; Li, X.; Hratchian, H. P.; Izmaylov, A. F.; Bloino, J.; Zheng, G.; Sonnenberg, J. L.; Hada, M.; Ehara, M.; Toyota, K.; Fukuda, R.; Hasegawa, J.; Ishida, M.; Nakajima, T.; Honda, Y.; Kitao, O.; Nakai, H.; Vreven, T.; Montgomery, Jr., J. A.; Peralta, J. E.; Ogliaro, F.; Bearpark, M.; Heyd, J. J.; Brothers, E.; Kudin, K. N.; Staroverov, V. N.; Kobayashi, R.; Normand, J.; Raghavachari, K.; Rendell, A.; Burant, J. C.; Iyengar, S. S.; Tomasi, J.; Cossi, M.; Rega, N.; Millam, J. M.; Klene, M.; Knox, J. E.; Cross, J. B.; Bakken, V.; Adamo, C.; Jaramillo, J.; Gomperts, R.; Stratmann, R. E.; Yazyev, O.; Austin, A. J.; Cammi, R.; Pomelli, C.; Ochterski, J. W.; Martin, R. L.; Morokuma, K.; Zakrzewski, V. G.; Voth, G. A.; Salvador, P.; Dannenberg, J. J.; Dapprich, S.; Daniels, A. D.; Farkas, Ö.; J. B.; Ortiz, J. V.; Cioslowski, J.; Fox, D. J. Gaussian, Inc., Wallingford CT, 2016.
4. S. Wang, Y. Fan, D. Li, C. Sun, Z. Lei, L. Lu, T. Wang and F. Zhang, Nat. Commun., 2019, 10, 1058.
5. T. M. Beck and B. Breit, Eur. J. Org. Chem., 2016, 5839.
6. R. I. Zubatyuk, V. N. Baumer, A. L. Tatarets, L. D. Patsenker and O. V. Shishkin, Acta Cryst., 2004, E60, o2252.
7. X. Yao, W. Liu, W. Zhu, A. Tiemuer, F. Zhou, S. Yang, H. Yu, X. Qian and Y. Liu, Chem. Commun., 2020, 56, 8111.
